# The recombination landscape and multiple QTL mapping in a *Solanum tuberosum* cv. ‘Atlantic’-derived F_1_ population

**DOI:** 10.1101/2020.08.24.265397

**Authors:** Guilherme da Silva Pereira, Marcelo Mollinari, Mitchell J. Schumann, Mark E. Clough, Zhao-Bang Zeng, G. Craig Yencho

## Abstract

There are many challenges involved with the genetic analyses of autopolyploid species, such as the tetraploid potato, *Solanum tuberosum* (2*n* = 4*x* = 48). The development of new analytical methods has made it valuable to re-analyze an F1 population (*n* = 156) derived from a cross involving ‘Atlantic’, a widely grown chipping variety in the USA. A fully integrated genetic map with 4,285 single nucleotide polymorphisms, spanning 1,630 cM, was constructed with MAPpoly software. We observed that bivalent configurations were the most abundant ones (51.0∼72.4% depending on parent and linkage group), though multivalent configurations were also observed (2.2∼39.2%). Seven traits were evaluated over four years (2006-8 and 2014) and quantitative trait loci (QTL) mapping was carried out using QTLpoly software. Based on a multiple-QTL model approach, we detected 21 QTL for 15 out of 27 trait-year combination phenotypes. A hotspot on linkage group 5 was identified as QTL for maturity, plant yield, specific gravity and internal heat necrosis resistance over different years were co-located. Additional QTL for specific gravity and dry matter were detected with maturity-corrected phenotypes. Among the genes around QTL peaks, we found those on chromosome 5 that have been previously implicated in maturity (*StCDF1*) and tuber formation (*POTH1*). These analyses have the potential to provide insights into the biology and breeding of tetraploid potato and other autopolyploid species.

## 1 Introduction

Polyploids can be classified based on the meiotic pairing dynamics of their duplicated sets of chromosomes. Allopolyploids are formed by the genomes of more than one ancestor species, which generally implies in disomic inheritance, where bivalent and preferential chromosome pairing is observed. Autopolyploids are formed by the genome duplication of a single ancestor species, such that polysomic inheritance is expected, with both bivalent and multivalent random chromosome pairing (Comai, 2005; Zielinski and Scheid, 2012). In autopolyploids, multivalent formation is one of the conditions for double reduction, a phenomenon in which sister chromatid segments end up in the same gamete (Huang *et al*., 2019). In order to investigate these complicated inheritance patterns, genetic mapping studies in autopolyploid species still need to deal with the molecular marker system limitation. Even though up to four different alleles (e.g. *abcd*) are expected in a locus of an outcrossing, autotetraploid individual, biallelic single nucleotide polymorphisms (SNPs) can be characterized, at best, by the dosage of an arbitrary allele, say *A*, as nulliplex (*aaaa*), simplex (*Aaaa*), duplex (*AAaa*), triplex (*AAAa*) or quadriplex (*AAAA*).

Several polyploid species are agriculturally important (Bennett, 2004). Particularly, potato is the third most cultivated food crop in the world after rice and wheat, with more than 17.6 million hectares cultivated worldwide in 2018, and a production exceeding 368.2 million tons (FAO, 2020). Yet advancement in the breeding of new varieties is still relatively slow. ‘Russet Burbank’, which still accounts for a large amount of the acreage planted in the United States, was developed in the early 1900s (Bethke *et al*., 2014), and ‘Atlantic’, a major chipping variety grown in the country, was originally released in 1976 (Webb *et al*., 1978). One of the reasons of such relatively slow breeding process is the relative complexity of the potato genome. Cultivated potato, *Solanum tuberosum*, is a highly heterozygous autotetraploid (2*n* = 4*x* = 48) outcrossing crop, which makes it very difficult to combine potentially beneficial characteristics through conventional breeding methods. In fact, many agronomic traits are polygenic, and show continuous distributions (Ghislain and Douches, 2020), where genotype-by-environment is usually important and need to be assessed.

The genome complexity has also impacted on the correct assessment of dosage from molecular markers, impeding proper linkage map construction and quantitative trait loci (QTL) mapping. Research has been conducted over the years to develop molecular techniques to facilitate potato breeding. However, since the first molecular genetic maps of potato were published (reviewed by Mann *et al*., 2011), very few breeding programs have begun to employ marker-assisted selection routinely (Slater *et al*., 2014). Some of the exceptions to this are for genetically simpler traits such as potato cyst nematode resistance (Schultz *et al*., 2012) and potato virus Y (Hämäläinen *et al*., 1997) and other disease resistance loci (Ramakrishnan *et al*., 2015). Genomics-assisted breeding for more complex traits such as yield are still elusive, and more studies on the genetic architecture of these traits as well as on genome-wide prediction (Endelman *et al*., 2018) are needed.

The early linkage maps constructed for potato have incorporated qualitatively scored molecular markers (Mann *et al*., 2011). However, this traditional presence-absence based scoring limited not only the ability to explore the full range of segregating markers, but also to build integrated genetic maps for full-sib populations (Luo *et al*., 2004). Advances in molecular technologies have been made available to breeders with the publication of the diploid potato genome sequence (Xu *et al*., 2011) and the subsequent development and release of the Illumina Infinium^®^ 8,303 Potato Array (Felcher *et al*., 2012) through the USDA-NIFA Solanaceae Coordinated Agricultural Project (SolCAP). The original potato array, which was subsequently upgraded to a 12k form, was made up of 8,303 SNPs and has been shown to give very good coverage of the published potato genome (Felcher *et al*., 2012; Sharma *et al*., 2013). Utilizing Illumina’s technology of dual colored fluorescents to label the different nucleotides allows for scoring allele dosage (Schmitz Carley *et al*., 2017; Zych *et al*., 2019). Along with the development of an array that uses dosage-sensitive SNPs, new analytical methods have recently been developed to effectively utilize this information in reconstructing linkage maps (Hackett *et al*., 2014; Bourke *et al*., 2018; Mollinari and Garcia, 2019) and mapping QTL (Hackett *et al*., 2016; Chen *et al*., 2018) in autotetraploid species.

Novel linkage and QTL mapping approaches (Mollinari *et al*., 2020; Pereira *et al*., 2020) have enabled the exploration of the meiotic configurations and the detection of multiple QTL for traits of interest, offering breeding opportunities and insights into the biology of polyploid species. Particularly, Mollinari *et al*. (2020) presented a detailed characterization of the polysomic inheritance in an even more complex autopolyploid species, the hexaploid sweetpotato, *Ipomoea batatas* (2*n* = 6*x* = 90). In tetraploid potato, double reduction has been documented for several mapping populations (e.g. Bourke *et al*., 2015), but these studies lack a comprehensive assessment of multivalent formation and preferential chromosome pairing. As haplotypes and thus the patterns of genome transmission from parents to progeny had been inferred, Pereira *et al*. (2020) were able to estimate additive relationship based on identity-by-descent along the genetic map. Such an information was ultimately used in a random-effect model approach, facilitating multiple QTL mapping. In fact, single-QTL models used so far may have hindered the discovery of putative loci underlying the variation of agriculturally important traits in the species. Comparatively, multiple-QTL models have been shown to have increased detection power and to provide the statistical basis for separation of linked QTL (Pereira *et al*., 2020).

The advancement of molecular technologies and analytical methods, coupled with newly collected data, have made it valuable to re-analyze the B2721 potato mapping population, derived from the ‘Atlantic’ cultivar (McCord *et al*., 2011a, 2011b; Schumann *et al*., 2017). The aims of this study were (*i*) to perform SNP dosage calling and create a fully phased, integrated linkage map in order to study the meiotic configurations in the B2721 mapping population, and (*ii*) to carry out multiple QTL-based analyses for yield, foliage maturity, dry matter, specific gravity, skin texture and internal heat necrosis resistance evaluated over four years. This study has allowed us to infer the tetraploid inheritance mechanisms from a linkage analysis perspective, and to better understand these traits at the genetic level in order to bring breeders closer to marker-assisted selection.

## 2 Material and Methods

### 2.1 Mapping population and field experiment

The B2721 family consisted of 156 progenies and was created by crossing ‘Atlantic’ × B1829-5. This bi-parental mapping population was chosen based on its segregation for internal heat necrosis (IHN), a non-pathogenic physiological disorder characterized by brownish spots in the tuber parenchyma (Yencho *et al*., 2008). ‘Atlantic’ is susceptible to IHN, whereas B1829-5, an advanced round white clone from the USDA-ARS Beltsville potato breeding program, is resistant. The B2721 population also segregates for a wide range of agronomic traits of interest, and these were our focus in this work, although IHN related traits will also be considered. In this study, we used B2721 data previously collected in 2006, 2007 and 2008 (McCord *et al*., 2011b, 2011a) as well as additional data collected in 2014 (Schumann *et al*., 2017).

The field designs for years 2006, 2007 and 2008 were described by McCord *et al*. (2011a,b). Briefly, in 2006, the population was planted in an unreplicated trial with six plants per clone, whereas in 2007 and 2008, the population was planted in two replications with 10 plants per plot in a randomized complete block design, the same design also used in 2014. All experiments were carried out at the Tidewater Research Station (35°52’20” N, 76°39’33” W) in Plymouth, North Carolina, in a Portsmouth fine sandy loam. In 2014, untreated seed pieces were planted on March 21 and the harvest date was July 15 (a total of 116 days). Standard crop production methods used for potato cultivation in eastern North Carolina were used for all experiments. At harvest, plots were dug using a chain digger and harvested by hand. Tubers were then washed, culled for excessive rots and defects, and phenotype measurements were obtained.

### 2.2 Phenotypic data and analyses

Phenotyping for all IHN-related and agronomic traits in 2006, 2007 and 2008 was described previously (McCord *et al*., 2011b). In 2014, in addition to IHN incidence (NI) and IHN severity (NS) (Schumann *et al*., 2017), the B2721 population was evaluated for five other traits: plant yield (PY), foliage maturity (FM), dry matter (DM), specific gravity (SG) and skin texture (ST). PY was measured as total plot weight minus the tubers culled divided by 10 (number of plants per plot) in kg/plant. FM was measured as the area under the senescence progress curve (Shaner and Finney, 1977), based on foliage ratings taken on June 9, June 25 and July 2, using a scale of 0 (0% yellowing) to 5 (100% senesced) with half point increments. For DM determinations, tubers were quartered from stem end to bud end using an Easy Wedger, Model N55550-4 (NEMCO Inc., Hicksville OH). For each plot, one quarter from four different tubers was placed into plastic Whirl-Pak^®^ bags. Samples were then weighed, frozen at –20 °C, lyophilized, and weighed again. DM was calculated as the proportion between dry and fresh weights for each sample, in percentage. SG was calculated using the formula *SG* = *w*_air_/(*w*_air_ − *w*_water_) by using the air weight (*w*_air_) and water weight (*w*_water_) for each plot. ST was rated on a 0 (smooth) to 3 (russeted) scale. Finally, NI was calculated as the proportion of tubers showing any sign of IHN over all evaluated tubers, and NS was rated on a 9 (no IHN signs) to 1 (completely necrotic) scale and averaged out for all evaluated tubers (Schumann *et al*., 2017).

In order to estimate mean-basis broad-sense heritability (*H*^2^) (Holland *et al*., 2003), a joint analysis with data from 2007, 2008 and 2014 (replicated trials) was carried out, where years, blocks within years, genotypes and genotype-by-year interactions were treated as random effects. Except for 2006 data (single observation), adjusted means were obtained separately for each trait-year combination based on a mixed model with genotypes and blocks as fixed and random effects, respectively, plus the random residual error using ASReml-R v. 4.1.0 (Butler *et al*., 2018). The phenotype means were named after a trait abbreviation followed by two-digit year (trait-year combination, e.g. PY06) and later used for QTL mapping purposes. Progeny variation within each phenotype was explored with boxplots from R package ggplot2 (Wickham, 2016). Pearson’s correlations were computed using Rpackage psych v. 1.9.12 (Revelle, 2018), and graphical visualization through network and correlation plots were obtained using R packages corrr v. 0.4.0 (Ruiz *et al*., 2019) and corrplot v. 0.84 (Wei *et al*., 2017). As an approach to investigate genotype-by-environment (GE) interaction, genotype (G) plus GE interaction (GGE) biplot analysis (Yan and Kang, 2003) was carried out using R package GGEBiplots v. 0.1.1 (Dumble *et al*., 2017) based on the standardized means. All analyses were based on R software v. 3.6.0 (R Core Team, 2020).

### 2.3 Genotypic data, dosage calling and linkage map construction

Genotyping was performed using the Illumina Infinium^®^ 8,303 Potato Array (Felcher *et al*., 2012). The intensity of each fluorescent labeled nucleotide was read using the Illumina iScan Reader and imported into GenomeStudio. The ordered pair of normalized intensities (*x, Y*) from the two allelic variants was transformed into polar coordinates 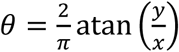 and 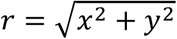 to proceed with the dosage calling of the parents and offspring using the R package ClusterCall (Schmitz Carley *et al*., 2017). Uninformative markers as well as those resulting in unclassified parents were filtered out resulting in 5,599 SNPs.

Allele dosage from these markers were imported into MAPpoly v. 0.1.0 (Mollinari and Garcia, 2019) together with their respective chromosome positions (*S. tuberosum* genome v. 4.0.3). After testing for segregation distortion (∼8.9 × 10^−6^ using Bonferroni correction) and filtering out redundant markers, the pairwise recombination fraction of the 4,812 SNPs (∼11.6 million pairs) were computed for all possible linkage phases and the most likely configuration was selected. Markers were assembled into 12 linkage groups (LGs) using the Unweighted Pair Group Method with Arithmetic Mean (UPGMA) algorithm. Since 91.2% of markers within the groups coincided with their respective chromosomes, groups were formed using exclusively the genome information. Markers were ordered according to the reference genome (Xu *et al*., 2011) and phased using function ‘est_rf_hmm_sequential’. Briefly, this function sequentially inserts markers into the map using two-point information to eliminate unlikely linkage phases (logarithm of the odds, LOD < 10.0), and for the remaining configurations, the multipoint LOD score is used (LOD < 10.0). If the insertion of a marker still results in less than 60 linkage phase configurations, a next marker was inserted and tested in the multiple configurations until the map is completed. In the final step, the multipoint recombination fraction was estimated allowing a global genotyping error of 5%.

Next, the conditional probabilities of the 36 possible genotypes were obtained for every cM of the linkage map for all individuals in the offspring. These probabilities were used to assess the meiotic process that formed the offspring and for further QTL analysis. Probabilistic profiles for pairing behavior were obtained for all LGs for the three possible meiotic pairing configurations, i.e., *ab*/*cd, ac*/*bd*, and *ad*/*bc*, where *a, b, c* and *d* denote the homologs in parent ‘Atlantic’. The notation *ab*/*cd* indicates that homolog *a* paired with *b*, and *c* paired with *d*, for example. The same reasoning applies to parent B1829-5 with homologs *e, f, g* and *h*. Also, the probabilistic haplotypes of all individuals in the offspring were reconstructed and the crossing-over points and respective homologs involved in the exchange were inferred. Using this information and the heuristic algorithm presented by Mollinari *et al*. (2020), recombination chains were assembled for all offspring individuals, and the homologs involved in each meiosis were assessed. Recombination chains with more than two homologs involved imply that a multivalent formation was present during the meiosis.

### 2.4 QTL mapping and gene search

The conditional genotype probabilities calculated from the inferred genetic map were used to a compute sib-pair additive relationship based on identity-by-descent for every cM position in the map. Using the R package QTLpoly v. 0.2.0 (Pereira *et al*., 2020), we detected putative QTL positions using the random-effect multiple interval mapping (REMIM) model for each phenotype (trait-year combination) based on a forward-backward procedure. First, putative QTL were consecutively added to the model based on a forward search with a relatively relaxed genome-wide significance threshold (*α* = 0.20). Then, a backward elimination step was carried out under a more stringent threshold (*α* = 0.05). The variance components associated with the putative QTL were tested using linear score statistics (Qu *et al*., 2013), whose associated *P*-values were used in QTL profile visualization with the logarithm of *P*-values [*LOP* = − log_10_(*P*)]. The genome-wide significance was assessed using a score-based resampling method (Zou *et al*., 2004). A final multiple QTL model was fitted with the selected positions, and QTL genotype best linear unbiased predictions (BLUPs) were used to compute additive allele effects (Kempthorne, 1955). QTL heritability 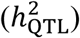 was calculated as the ratio between the variance associated with the QTL and the total variance.

In order to compare different approaches, we also ran the fixed-effect interval mapping (FEIM) model (Hackett *et al*., 2014) as implemented in the QTLpoly package (Pereira *et al*., 2020). Using the same conditional genotype probabilities, a single-QTL model was used to fit six additive effects (as one additive effect from each parent is taken as reference) at every cM position. The model with a fitted QTL was compared to a null model (with no QTL) using likelihood ratio tests (LRT). LRT statistics were converted into LOD scores, and QTL were declared when the LOD score reached a threshold (*α* = 0.05) based on 1,000 permutation tests (Churchill and Doerge, 1994). In both REMIM and FEIM analyses, a window size of 20 cM was used to avoid that two closely linked positions were declared as QTL. Approximate 95% QTL support intervals were estimated by dropping 1.5 from the QTL peak *LOP* (REMIM) or *LOD* (FEIM) (Pereira *et al*., 2020).

Based on the QTL peaks from REMIM, we searched for candidate genes in the region delimited by markers on the left and on the right of the QTL peak or within 200 kbp each side from the QTL peak, whichever was the largest, on the *S. tuberosum* v. 4.03 genome (ST4.03) using the PhytoMine tool (https://phytozome.jgi.doe.gov/phytomine/) by Phytozome v. 12 (Goodstein *et al*., 2012). Finally, we used the Web Gene Ontology Annotation Plot (WEGO) v. 2.0 (Ye *et al*., 2018; http://wego.genomics.org.cn/) to count and plot classifications of Gene Ontology (GO) terms for annotated genes within our QTL regions.

## 3 Results

### 3.1 Trait correlations and GE interactions

Based on the phenotype means (see Supplementary File S1), variation was found for all evaluated traits over the four years of evaluation (Figure 1A), indicating that they were segregating in the B2721 population. As the parental means were within the population phenotypic range, we could observe transgressive segregation for all trait-year combinations. PY06 has shown quite distinct median and broader variation in comparison to the following years, which is likely because there were no replicates in 2006, so that the environmental error could not be modeled. In contrast, FM14 exhibited a narrower variation when compared to the previous years. This was rather expected, since in 2014 maturity evaluation was carried out in a shorter time-window (24 days) in comparison to 2007 and 2008 (30 days). The overall change in ST14 scores when compared to the previous years was likely due to differences in evaluator scoring. Individuals seemed to be less affected by IHN in 2008, as NS08 and NI08 have shown distinct variation and median in comparison to the other years. Broad-sense heritability (*H*^2^) ranged from 0.39 (DM) to 0.81 (SG), with similarly high values for PY (0.78), FM (0.70) and ST (0.79). For IHN-related traits, NS (0.74) showed a relatively higher heritability when compared to NI (0.66).

**Figure 1.**
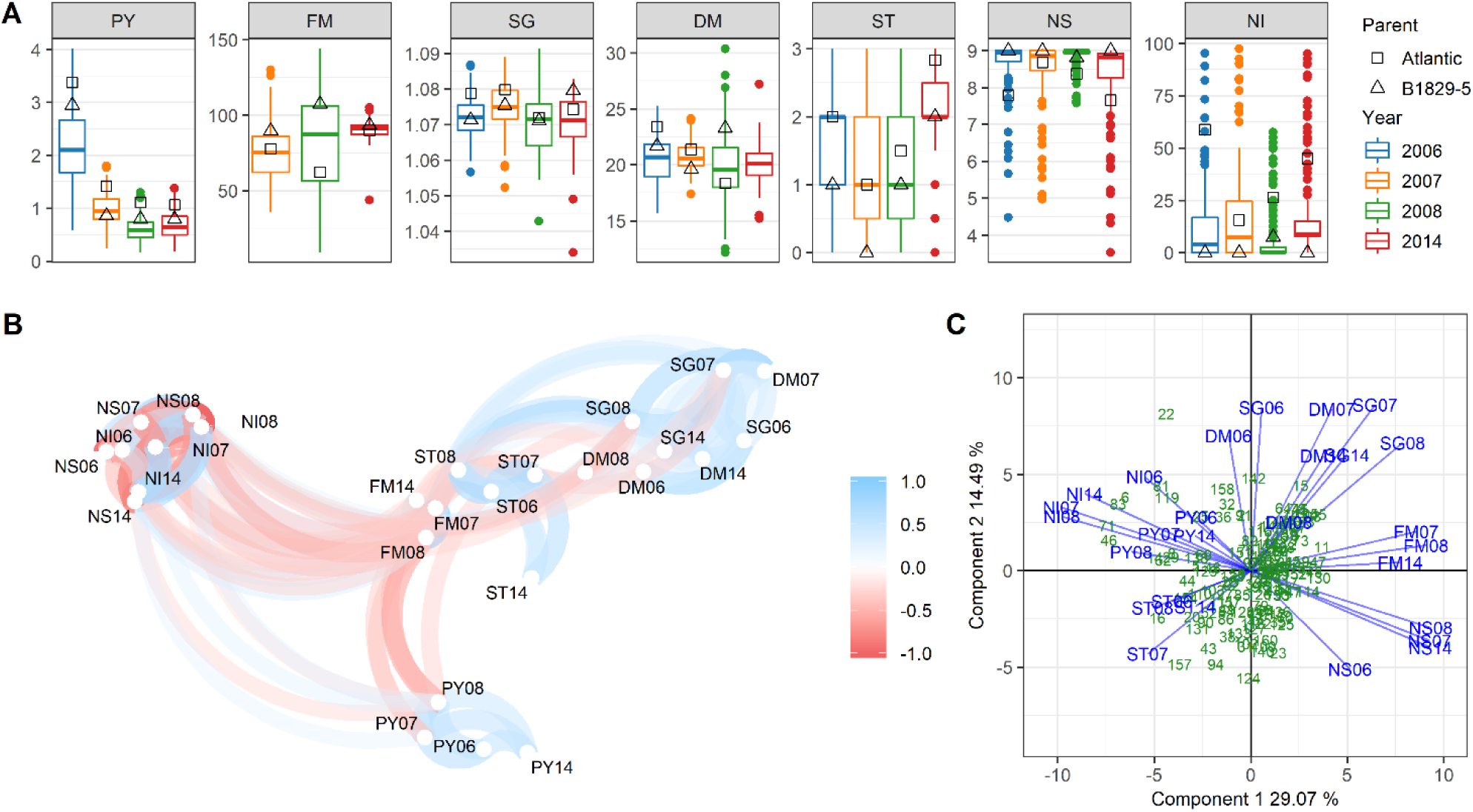
B2721 mapping population phenotypes evaluated for four years (2006-8 and 2014). (A) Boxplots showing distribution of full-sib means along with parental means. (B) Network plot showing positive (blue) and negative (red) significant correlations (|*r*| ≥ 0.26, *P* < 0.001). (C) GGE biplot showing the distribution of full-sibs (green) and phenotypes (blue) on the two first principal components. Traits: plant yield (PY), foliage maturity (FM), specific gravity (SG), dry matter (DM), skin texture (ST), and internal heat necrosis severity (NS) and intensity (NI). The phenotypes were named after a trait abbreviation followed by two-digit year (e.g. PY06).

The correlation among variables was explored using a network plot (Figure 1B), where path color and transparency represent the signal and strength of the correlation, respectively. The variable positioning is given by multidimensional scaling that leverages the magnitude of the correlations, i.e. the closer the variables, the higher the absolute values of their correlations (Ruiz *et al*., 2019). In general, the phenotype of the same trait evaluated in different years clustered together. As expected, NS was close to NI as well as DM to SG, thus indicating high correlation, because each one of these trait pairs are different ways of assessing the same characteristic (IHN response and solids content, respectively). FM appeared centrally in this plot as it was correlated with most variables either negatively (e.g. with PY and NI) or positively (e.g. with SG, DM and NS). Note that for this network plot, a subset of 144 full-sibs with complete observations among all traits was ultimately used. For correlation estimates based on pairwise complete observations, please see Supplementary Figure S1 and Supplementary File S2. Even though data from 2006 was originated from a single replication, they were in general agreement with the adjusted means from the other years. DM observations were the least correlated ones across years in comparison to all the other trait sets (see Supplementary Figure S1).

GGE biplot was used to investigate the existence of GE interaction (Figure 1C), where each year was considered an environment. The distribution of full-sibs along the first two principal components, which accounted for 43% of the variation, did not show any clustering. This was expected as mapping populations usually consists of non-selected individuals. However, some full-sibs were found to be more scattered in the top-left (fourth) quadrant, indicating the most affected individuals by IHN (i.e. high NI). In contrast, more individuals were found to be less scattered in the bottom-right (second) quadrant, where NS vectors lie, indicating smaller differentiation when full-sibs were less susceptible to IHN. Relatively shorter vector lengths of PY, ST and DM08 indicated limited capacity to discriminate genotypes using these traits, in comparison to longer vectors. Only relatively weak GE interaction was observed, since phenotypes from different years consistently exhibited the same vector orientation for each trait, with small angles between vectors. Greater angles between vectors of phenotypes from 2006 (such as DM06, SG06 and NS06) and from the remaining years should be treated carefully, as these 2006 measurements were based on single plots. The same vector orientation for SG and DM indicated that these traits were positively correlated, whereas the opposite orientations for NS and NI vectors revealed their negative correlations.

### 3.2 Linkage analysis and inheritance mechanisms

The normalized intensities from the Illumina Infinium^®^ 8,303 Potato Array were obtained for the B2721 population (see Supplementary File S3). The dosage calling procedure yielded 5,599 informative SNPs with 1.05% of missing data. Fourteen percent of the SNPs were filtered out due to segregation distortion and redundancy resulting in 4,812 informative markers: 1,311 (27.3%) simplex, 1,234 (25.6%) double-simplex, and 2,267 (47.1%) multiplex (i.e. from duplex to quadriplex; see Supplementary Figure S2). The complete linkage map (see Supplementary Table S1, Supplementary Figure S3 and Supplementary File S5) consisted of 12 LGs with eight haplotypes each, with four homologs per parent. A total of 4,285 markers spanned 1,629.99 cM in length, with an average density of 2.64 SNPs/cM. Linkage group lengths ranged from 106.20 cM (LG 5) to 205.88 cM (LG 1), with average LG length of 135.83 cM. The scatterplots of the physical distance in *S. tuberosum* genome v. 4.03 versus genetic distance in the 12 LGs in B2721 population map is shown in Supplementary Figure S4.

The probabilistic pairing profiles showed no preferential pairing between homologs in both parents (see Supplementary Figure S5). Among all meiotic configurations, only 7.3% and 9.8% were inconclusive for parents ‘Atlantic’ and B1829-5, respectively. From the remaining configurations (Figure 2), the percentage of cases with no crossing-over varied from 6.8% (LG 1, parent ‘Atlantic’) to 40.7% (LG 11, parent B1829-5), with 24.2% on average. Configuration involving two chromosomes with at least one crossing-over (i.e. bivalent configurations), were the most abundant varying from 51.0% (LG 11, parent B1829-5) to 72.4% (LG 9, parent ‘Atlantic’), with a mean of 62.3%. Multivalent configurations, i.e., involving three or four homologs, ranged from 2.2% (LG 8, parent B1829-5) to 39.2% (LG 1, parent ‘Atlantic’), with 13.5% on average.

**Figure 2.**
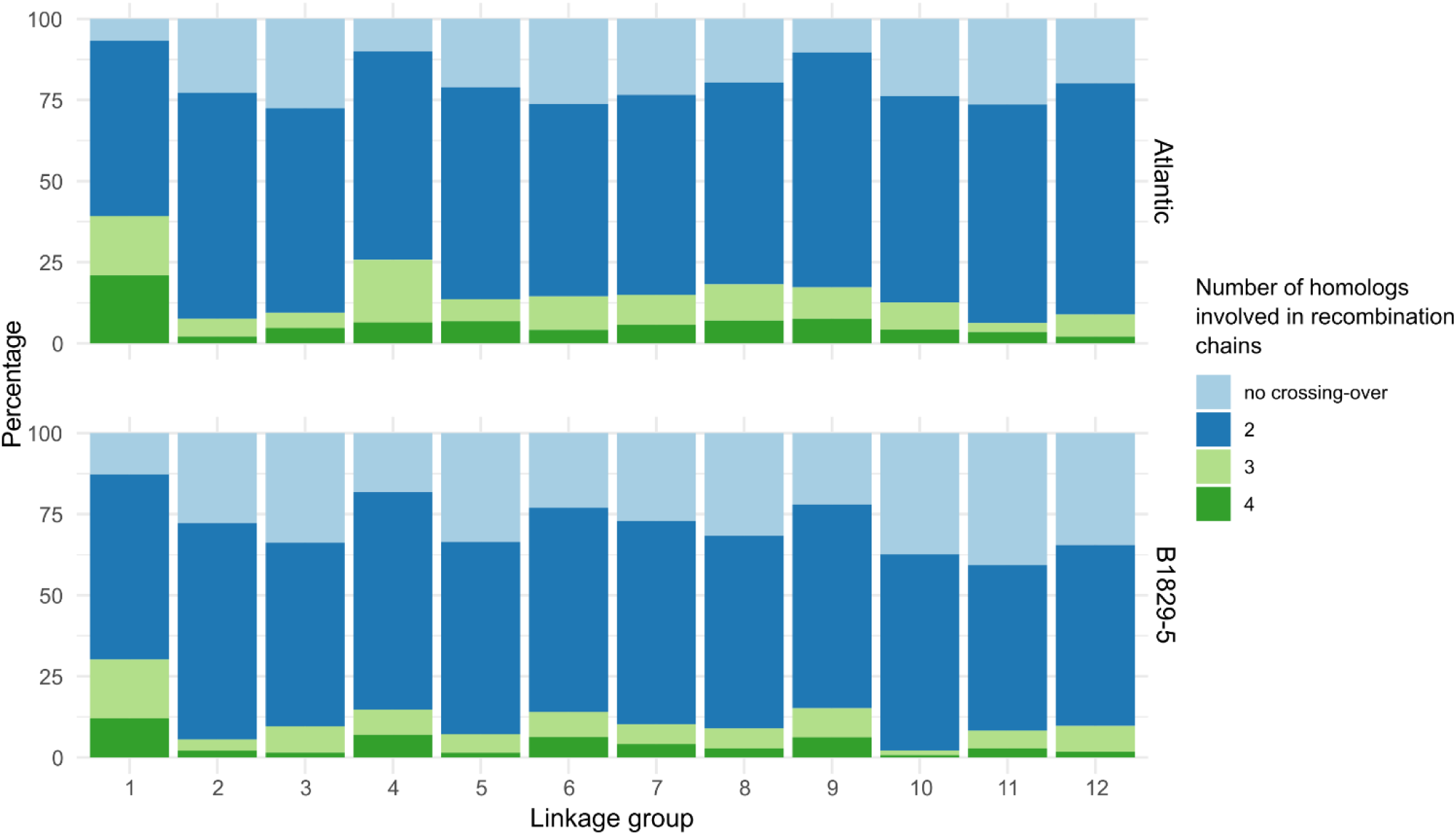
Distribution of number of homologs involved in a recombination chain.

### 3.3 Multiple QTL mapping and candidate genes

From the REMIM analysis, a total of 21 QTL were mapped in 15 out of 27 evaluated trait-year combinations (Table 1; see Supplementary Figure S6). One QTL was detected for 10 phenotypes each (for PY06, PY08, SG07, SG08, ST07, ST08, NS06, NS08, NI06 and NI08). The other five phenotypes have shown two (for FM07, FM14, SG06 and NI07) or three (for FM08) QTL each. No QTL were mapped for 12 phenotypes (PY07, PY14, SG14, DM06, DM07, DM08, DM14, ST06, ST14, NS07, NS14 and NI14), mostly from years 2006 and 2014. Ten QTL were mapped on LG 5, four on LG 1, three on LG 7, and one on LGs 2, 3, 4 and 9 each. QTL heritability 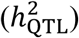 ranged from 7.1% (FM08 on LG 7) to 64.0% (FM08 on LG 5), including five QTL with higher 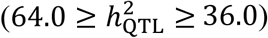, 10 with moderate 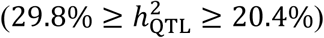, and the remaining six QTL with lower 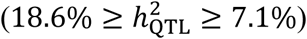 heritability (Table 1; Figure 3). In comparison to REMIM results, FEIM detected 17 QTL from 12 phenotypes (see Supplementary Figure S8). On the one hand, FEIM did not identified QTL previously identified for PY06 (on LG 5), FM08 (on LGs 1 and 7), SG06 (on LG 3), NI06 (on LG 1) and NI07 (on LGs 1 and 5). On the other hand, FEIM identified additional QTL for SG06 (on LGs 8 and 10), ST08 (LG 9), NI14 (on LG 5) (see Supplementary Table S). The proportion of variance explained (PVE) by FEIM-derived QTL ranged from 13.0% to 52.7%.

**Table 1.** Random-effect multiple interval mapping (REMIM) for B2721 traits evaluated for four years (2006-8 and 2014).

| Trait<sup>a</sup> | QTL | Linkage group | Position (cM) | SI (cM)<sup>b</sup> | Score | P-value | $\sigma^2_{\text{QTL}}$<sup>c</sup> | $h^2_{\text{QTL}}$<sup>d</sup> (%) |
| --- | --- | --- | --- | --- | --- | --- | --- | --- |
| PY06 | 1 | 5 | 32 | 13-42 | 137.41 | 4.70e-05 | 1.11e-01 | 20.4 |
| PY08 | 1 | 5 | 46 | 0-53 | 168.95 | 7.21e-06 | 2.17e-02 | 36.0 |
| FM07 | 1 | 5 | 26 | 18-29 | 355.99 | <2.22e-16 | 3.36e+02 | 57.7 |
|  | 2 | 7 | 66 | 52-70 | 167.43 | 8.98e-07 | 7.28e+01 | 12.5 |
| FM08 | 1 | 1 | 9 | 0-32 | 101.58 | 2.16e-04 | 1.17e+02 | 7.4 |
|  | 2 | 5 | 27 | 0-41 | 425.04 | <2.22e-16 | 1.01e+03 | 64.0 |
|  | 3 | 7 | 65 | 35-71 | 104.85 | 2.00e-04 | 1.13e+02 | 7.1 |
| FM14 | 1 | 5 | 18 | 0-29 | 197.01 | 9.77e-08 | 7.12e+00 | 29.8 |
|  | 2 | 7 | 66 | 54-70 | 138.45 | 1.57e-05 | 4.01e+00 | 16.8 |
| SG06 | 1 | 2 | 97 | 34-117 | 125.63 | 6.65e-05 | 6.69e-06 | 15.9 |
|  | 2 | 3 | 44 | 31-50 | 109.89 | 1.49e-04 | 8.94e-06 | 21.2 |
| SG07 | 1 | 5 | 19 | 4-32 | 135.93 | 3.26e-05 | 1.39e-05 | 27.6 |
| SG08 | 1 | 5 | 28 | 15-32 | 240.07 | 1.47e-08 | 4.24e-05 | 48.9 |
| *SG07 | 1 | 2 | 104 | 68-125 | 131.20 | 8.32e-05 | 8.35e-06 | 23.0 |
| *SG08 | 1 | 2 | 115 | 60-124 | 143.00 | 5.49e-05 | 7.80e-06 | 20.9 |
| *DM07 | 1 | 2 | 115 | 63-121 | 128.17 | 1.40e-04 | 2.62e-01 | 14.9 |
|  | 2 | 6 | 36 | 31-73 | 134.38 | 3.58e-05 | 4.13e-01 | 23.3 |
| ST07 | 1 | 9 | 41 | 27-82 | 109.68 | 1.97e-04 | 1.59e-01 | 21.9 |
| ST08 | 1 | 4 | 2 | 0-4 | 234.05 | 1.08e-08 | 6.32e-01 | 48.1 |
| *ST08 | 1 | 4 | 2 | 0-4 | 255.84 | 3.62e-09 | 5.45e-01 | 46.9 |
| NS06 | 1 | 1 | 0 | 0-52 | 113.07 | 1.29e-04 | 1.17e-01 | 22.9 |
| NS08 | 1 | 5 | 26 | 12-32 | 117.77 | 1.77e-04 | 2.84e-02 | 23.4 |
| NI06 | 1 | 1 | 0 | 0-69 | 110.07 | 1.66e-04 | 9.28e+01 | 22.0 |
| NI07 | 1 | 1 | 0 | 0-8 | 103.32 | 2.26e-04 | 1.39e+02 | 16.8 |
|  | 2 | 5 | 25 | 8-34 | 134.43 | 3.35e-05 | 1.99e+02 | 24.2 |
| NI08 | 1 | 5 | 27 | 12-32 | 131.28 | 6.54e-05 | 5.65e+01 | 28.5 |

\*Maturity-corrected phenotypes.
<sup>a</sup>Traits: plant yield (PY), foliage maturity (FM), specific gravity (SG), dry matter (DM), skin texture (ST), and internal heat necrosis severity (NS) and intensity (NI).
<sup>b</sup>~95% support interval (SI) based on $LOP - 1.5$ , in centiMorgans.
<sup>c</sup>Variance component associated with the QTL ( $\sigma^2_{\text{QTL}}$ ).
<sup>d</sup>QTL heritability ( $h^2_{\text{QTL}}$ ), in percentage.

**Figure 3.**
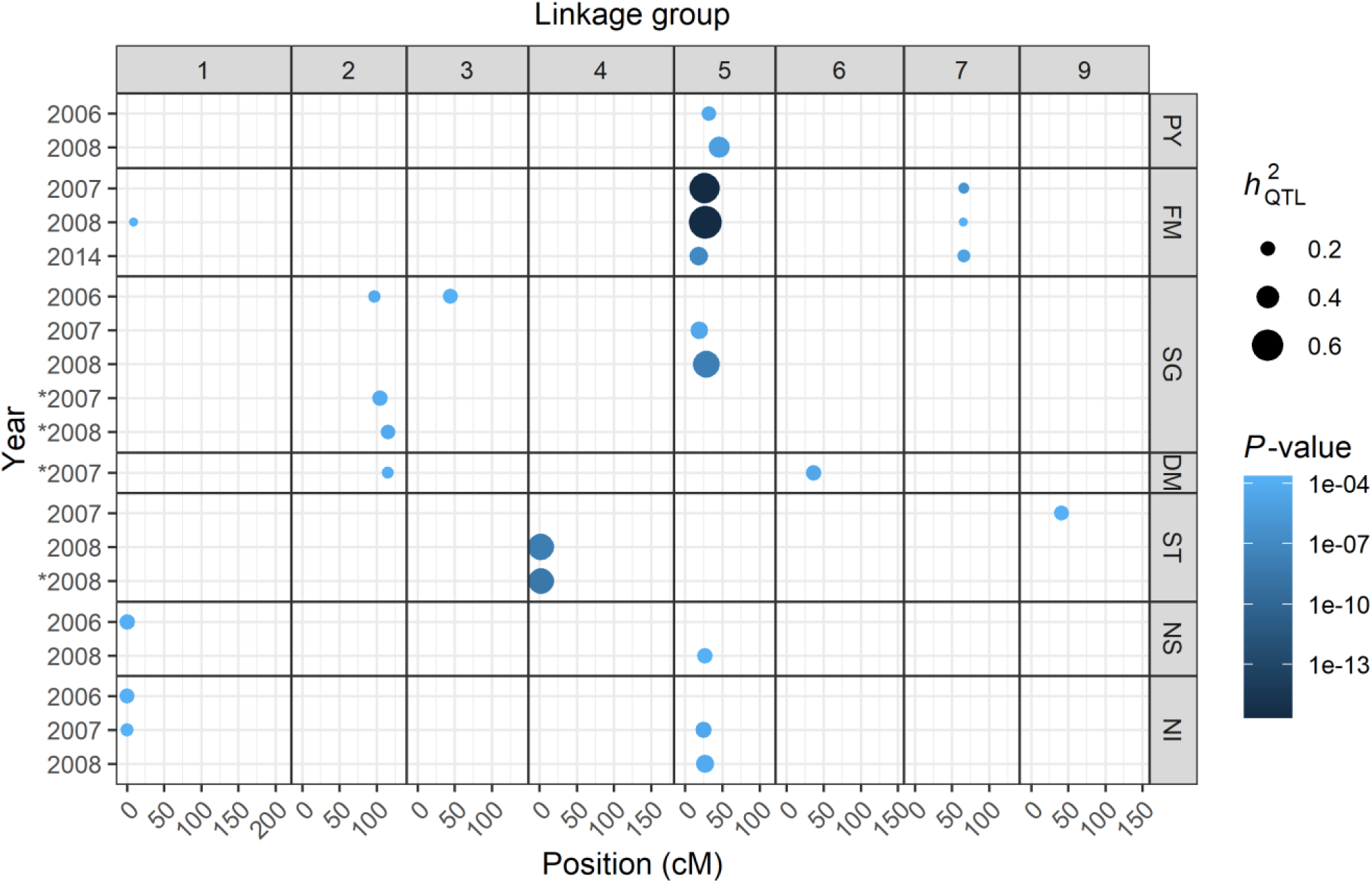
QTL identified for seven traits evaluated for four years (2006-8 and 2014). Dots are plotted according to QTL peak location along the linkage groups. Color scale represents *P*-values, and sizes are proportional to QTL heritability 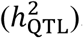. Traits: plant yield (PY), foliage maturity (FM), specific gravity (SG), dry matter (DM), skin texture (ST), and internal heat necrosis severity (NS) and intensity (NI). * indicates maturity-corrected phenotypes.

With 10 QTL on the proximal half of LG 5, this region was characterized as a QTL hotspot. The major QTL on LG 5 for FM was detected for all three years of evaluation (2007-8 and 2014). However, *LOP* profiles (see Supplementary Figure S6) showed suggestive QTL (at a lower *LOP* that did not reach the threshold), for instance, on LG 5 for PY06, SG14 and NS14 and NI14. For the five traits with QTL on LG 5 (PY, FM, SG, NS and NI), most QTL peaks were found at 29 cM and support intervals ranged from 0 to 50 cM. Figure 4 depicts additive allele effects of co-located QTL on LG 5 for these different traits evaluated in 2008. It is interesting to note that for the QTL sets PY08.1/NI08.1 and FM08.2/SG08.1/NS08.1, major allele effects appeared in the same direction within sets, but in opposite directions between them. As these QTL on LG 5 were responsible for explaining significant portion of the variation for these traits (Table 1), the QTL-based predicted means were highly correlated, either positively (PY with NI, and FM with SG and NS) or negatively (PY/NI with FM/SG/NS) (see Supplementary Figure S11). It is worth mentioning that selecting towards two out of four alleles per parent will tend to favor different sets of phenotypes depending on the alleles. For example, selecting allele pair *ac* from ‘Atlantic’ and *eg* from B1829-5 will likely increase PY and NI and decrease FM, SG and NS. Additive effects for all QTL can be found on Supplementary File S6.

**Figure 4.**
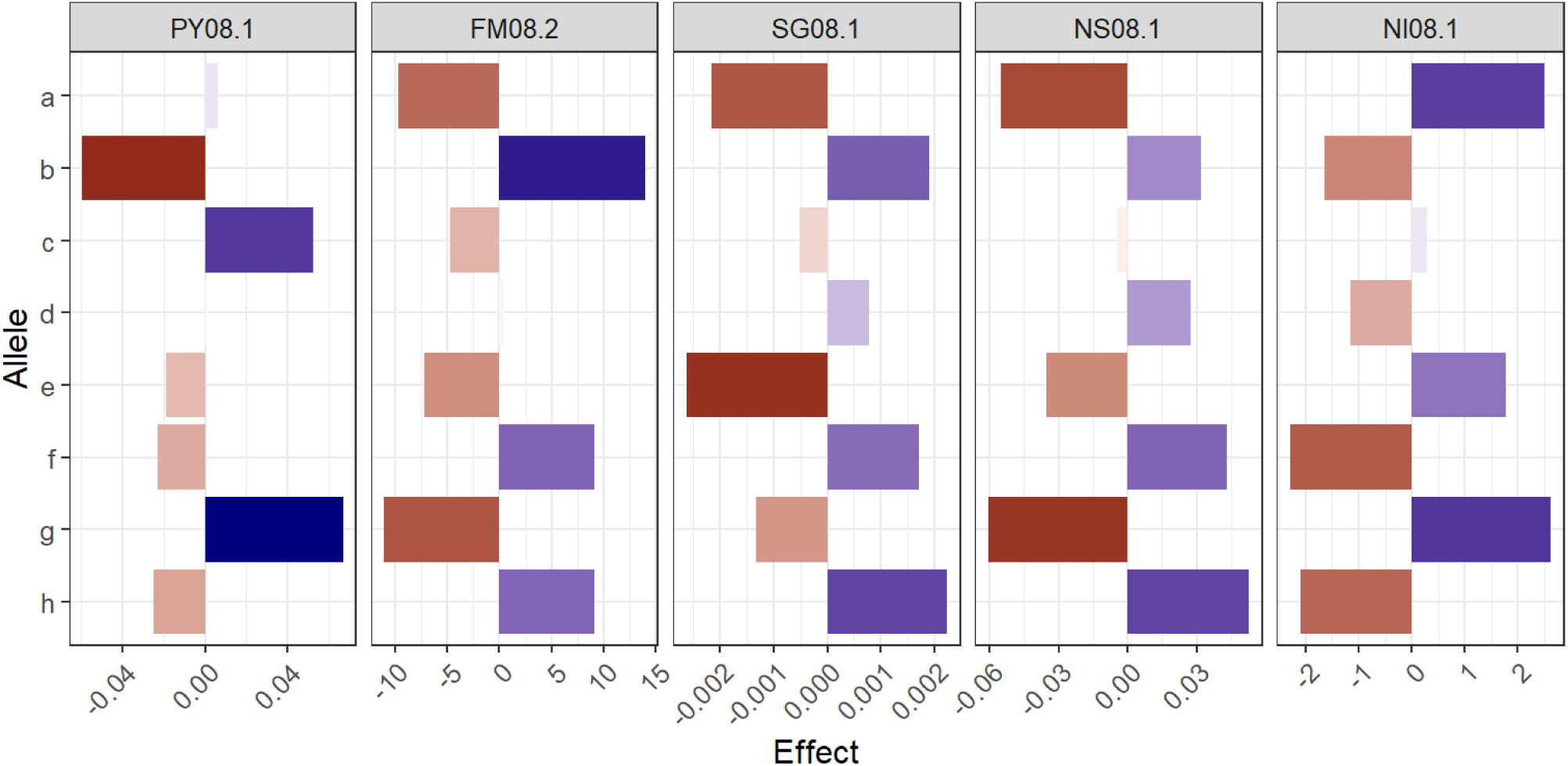
Additive allele effects for co-located QTL on LG 5 for five traits evaluated in 2008. Letters represent each parental haplotype from the linkage map (‘Atlantic’ = *a* through *d*, B1829-5 = *e* through *h*). Traits: plant yield (PY), foliage maturity (FM), specific gravity (SG), and internal heat necrosis severity (NS) and incidence (NI).

Due to such a large QTL effect on LG 5 for FM and its consistent impact on most traits, we fitted simple linear regressions of the form ***y*** = **1***β*_0_ + *β*_1_***x*** + ***e*** using the lm() function of R (R Core Team, 2020), with FM as the explanatory variable ***x*** and any other phenotype as dependent variable ***y*** from each respective year where FM was measured (i.e. 2007, 2008 and 2014). The vector of residuals ***e*** were used as maturity-corrected phenotypes (i.e. virtually free of FM influence; see Supplementary Figure S10) for new QTL mapping runs, as suggested by Bradshaw *et al*. (2004). We found that the slopes *β*_1_ were highly significant for all traits (\*\*\**P* < 0.001), except DM14 (\**P* < 0.05), and DM08, PY14, SG14 and ST14 (*P* > 0.1) (see Supplementary Table S2). REMIM analyses on maturity-corrected phenotypes resulted in five QTL (see Supplementary Fig. S7). A QTL for ST08 co-located with the one previously identified on LG 4. Other four QTL were newly identified: one QTL from each SG07, SG08 and DM07 were closely located on LG 2, and another QTL for DM07 was detected on LG 6 (Table 1; Figure 3). FEIM analyses returned eight QTL, from which two were previously identified for ST08 (LGs 4 and 9), and other six were newly detected for SG07 (LG 2), SG 08 (LGs 2 and 5), DM07 (LG 6), DM08 (LG 11), and NI14 (LG 4) (see Supplementary Table S4 and Supplementary Figure S9). As the QTL support intervals comprised too large of a region to manually curate (>13 Mb per QTL, on average), we focused our candidate gene search on the region within the markers flanking the QTL peak (∼400 kb, on average) (see Supplementary Table S3). A total of 533 annotated genes on *S. tuberosum* v. 4.03 genome were within these QTL regions, with 342 genes assigned to at least one GO term (see Supplementary File S7 and Supplementary Figure S12).

## 4 Discussion

### 4.1 The recombination landscape of B2721 mapping population

The ability to assess the intricate recombination landscape of an outcrossing, autopolyploid species encompass the construction of a highly informative map. That is, the parental maps have to be integrated between the two parent and must differentiate each one of the parental haplotypes. Two genetic maps have been built previously for the B2721 population. The first map used amplified length polymorphisms (AFLP) and a few simple sequence repeat (SSR) markers, and consisted of two separate linkage maps, one for each parent, with limited integration within parents (McCord *et al*., 2011b). The second map used dosage-sensitive SNP array, which led to full integration and haplotyping in the B2721 population (Schumann *et al*., 2017). However, this map contained 3,427 SNPs and spanned 1,397.86 cM, which is a relatively smaller but less saturated than our current map with 4,285 SNPs distributed along 1,629.99 cM. The ability to incorporate more markers with MAPpoly was possibly due to (*i*) the multipoint-based algorithm that considers all parental homologs and marker dosages simultaneously, and (*ii*) the genome-assisted approach for marker ordering. Schumann *et al*. (2017) used the software TetraploidSNPMap (Hackett *et al*., 2016), which relies on (*i*) simplex markers to assemble homologs, bridging these homologs afterwards using multidose markers, and (*ii*) two-point genetic distances to obtain the marker order by using the multidimensional scaling (MDS) algorithm (Preedy and Hackett, 2016). Increased marker saturation provides better support to estimate the conditional probabilities, which are used to infer pairing profiles during meiosis in a segregating population and to map QTL.

The probabilistic pairing profiles showed no preferential pairing between homologs in both parents (see Supplementary Figure S5), likely due to the cultivated potato’s autotetraploid nature. Consequently, alleles present in different homologs in the same homology group had an equal chance to recombine with alleles from other loci, amplifying the range of possible genotypes when compared to diploids or allotetraploids. In addition, most of the parental meiotic configurations based on the B2721 population were inferred as bivalents (62.3% on average) since they involved two homologs exchanging segments during the metaphase I (Figure 2). These results are in accordance with recent findings published by Choudhary *et al*. (2020). These authors presented a map of meiotic stages in *S. tuberosum* in one diploid and three tetraploid varieties using fluorescence *in situ* hybridization (FISH). They concluded that bivalent chromosome associations are the most common in metaphase I of tetraploid potatoes. Likewise, the broad range of multivalent signature rates observed in our study (2.2∼39.2%) was also observed by Choudhary *et al*. (2020) (7∼48%). Similarly, as their observations differed depending on variety and chromosome, in our study, the variation occurred by parents and by linkage groups. Thus, our meiotic assessment, although not as precise as a cytological analysis and prone to sampling errors, can be very helpful, serving as a proxy to evaluate meiotic configurations by using a straightforward expansion of linkage analysis.

### 4.2 Multiple QTL mapping for yield, quality and IHN related traits

Single-QTL models were previously applied to the first two linkage maps of B2721. These models were based on an interval mapping model, and the 95^th^ percentile (α = 0.05) of maximum LOD scores from 1,000 permutation tests per phenotype was used to declare QTL, with several additional suggestive QTL (below this threshold) also being recorded (McCord *et al*., 2011b, 2011a). One of the issues with the lack of integration within and between parental maps is the same region being declared as a QTL in separate parental haplotypes. For instance, for plant yield, McCord *et al*. (2011b) listed a total of seven QTL across three years, PY06 (one), PY07 (five) and PY08 (two), from both parental maps, in addition to 11 suggestive ones. Our multiple-QTL analyses resulted in a single co-located QTL region on LG 5 for each PY06 and PY08 (Table 1; Figure 3), with suggestive QTL for PY07 and PY14 (see Supplementary Figure S6). QTL on LG 5 for tuber yield have also been reported in both diploid (DM 1-3 516 R44 × RH89-039-16, *n* = 96) (Manrique-Carpintero *et al*., 2015) and tetraploid (‘Liberator’ × W4013-1,*n* = 110) (Rak *et al*., 2017) potato mapping populations.

For foliage maturity, McCord *et al*. (2011b) and Schumann *et al*. (2017) described major QTL for years 2007-8 on LG 5 as well as the one on LG 7. In the current study, another QTL on LG 1 was identified for FM08, likely due to increased detection power of REMIM. Despite the differences regarding phenotypic distribution for FM14 (Figure 1A), we were able to find QTL on LGs 5 and 7 (Figure 3). This LG 5 region is well known as a main source of QTL in potato. Prior to the release of the SNP potato array, a major QTL for plant maturity had already been reported on chromosome 5, also linked to a late blight (*Phytophthora infestans*) resistance locus (reviewed by Danan *et al*., 2011). More recently, additional studies noticed that the same region on LG 5 also underlies the variation of several tuber quality-related traits in addition to resistance to early blight (*Altenaria tenuis*) and Verticillium wilt (*Verticillium* spp.) (Massa *et al*., 2018), and of tuber-Cd and Zn concentration (Mengist *et al*., 2018), for example. For IHN-related traits, three QTL were identified on LG 5, one for each NS08, NI07 and NI08, using REMIM. FEIM identified an additional significant QTL on LG 5 for NI14 (see Supplementary Table S4 and Supplementary Figure S8). Previously, IHN-related QTL had been reported as suggestive only (Schumann *et al*., 2017), and this difference in significance is likely due to an improved map construction.

A major QTL on LG 5 was also found to co-locate for SG07 and SG08, while it appeared only as suggestive for SG14 and with no evidence for SG06, which in turn showed one QTL on each LGs 2 and 3. The evaluation of maturity-corrected phenotypes helped to detect this co-located QTL for DM07, and SG07 and SG08 on LG 2. An alternative to perform QTL discovery free of maturity influence, though, would be to use materials that do not show much variation for this trait, i.e. crosses between either late or early maturity cultivars. Specific gravity and dry matter are supposedly related with solid content in the tuber. In fact, the correlation among their adjusted means were generally high (up to 0.81 between DM07 and SG07; see Supplementary Figure S11 and Supplementary File S2), but the only set of co-located QTL between any SG and DM measures was that one on LG 2. It is worth to mention that their broad-sense heritability values (*H*^2^) differed greatly (0.39 for DM and 0.81 for SG), so that the ability to detect QTL for DM may have been affected because smaller proportion of the total variance could be attributed to genetic variance. Other mapping populations have revealed QTL on LG 5 for DM (12601ab1 × ‘Stirling’, *n* = 227) (Bradshaw *et al*., 2008) and SG (Manrique-Carpintero *et al*., 2015), where somewhat distinct heritability values were found (0.81 for DM and 0.46 for SG).

Skin texture was a trait which diverged the most from the other traits regarding QTL locations, specifically for not showing enough statistical evidence for QTL on LG 5 (see Supplementary Figure S6). In diploid potatoes, skin texture is believed to be controlled by three loci (Jong, 1981). In our B2721 population, we detected two QTL (one for ST07 and ST08 each), in regions on LGs 4 and 9 where no other traits have shown evidence for QTL in our study. That is, the apparent strong correlation observed between ST07 and FM07 (*r* = −0.44, *P* < 0.001), for example, could not be attributed to genetics, as correlation between QTL-based predicted means for these two traits was close to zero (*r* = −0.07, *P* = 0.40; see Supplementary Figure S11).

When performing QTL discovery in outbred polyploid populations, in addition to genome complexity, one must keep two factors in mind. First, in F1 populations, the fact that two parental genotypes are contrasting does not directly imply that major QTL will be detected. For instance, considering a biallelic QTL, a cross between two contrasting parents, *QQQQ* × *qqqq*, would not segregate (i.e. 100% *QQqq*). In order to be detectable, alleles need to be contrasting within parent, e.g. *Qqqq* × *qqqq* or *Qqqq* × *Qqqq* in the simplest cases, resulting in segregations such as 50% *Qqqq* : 50% *qqqq* or 25% *QQqq* : 50% *Qqqq* : 25% *qqqq*, respectively. We speculate that the allele effects of undetectable QTL towards IHN resistance, for example, did not differ significatively within each parent. For the QTL on LG 5 for NI08, the allele contributions to the population mean (∼6.1%) were ±2.5% at most (Figure 4). In other words, these effects would not change the fact that most individuals would be less susceptible (NI < 10%) as resistance could have been already conferred by these undetectable QTL.

If there are segregating QTL alleles, another factor that influences detection is small population sizes that may not allow to separate signal from noise properly. Consequently, thresholds for declaring a QTL are usually required to be more stringent in order to avoid false positives. In general, tetraploid potato QTL mapping studies have been performed in relatively small population sizes (∼150 or less individuals). In these cases, even very low *p*-values and high QTL heritability estimates, such as those found for FM07, FM08 and SG08, should be treated carefully as they can be relatively biased due to sampling. Finally, reduced population sizes also limit the ability to test closely linked versus pleiotropy for co-located QTL across different traits or environments accurately.

### 4.3 Candidate genes within QTL regions

Among the candidate genes (see Supplementary File S7), GO terms that overlapped DNA and protein binding functions with regulation of biological process were of particular interest. In total, 19 genes were found to have the same two GO terms (GO:0003677 and GO:0006355) attributed to the *S. tuberosum* CYCLING DOF (DNA-binding with one finger) FACTOR 1, *StCDF1* (PGSC0003DMG400018408, ST4.03ch05:4539029..4541329), which has been described as the transcription factor underlying the major QTL for maturity on LG 5 (Kloosterman *et al*., 2013). The SolCAP markers around the QTL peak for FM08 (c2_22959 at 4,434,048bp and c2_23052 at 4,906,728 bp) were found to be closer to *StCDF1* than the flanking markers for FM07 and FM14 (c2_11829 at 4,041,510 bp and c2_22986 at 4,279,075 bp), and the marker mentioned by Massa *et al*. (2015) (c2_22986 at 4,105,142). The same LG 5 region also includes yield-related QTL, where we found the *Potato homeobox 1* gene, *POTH1* (PGSC0003DMG400013493, ST4.03ch05:9248372..9258054), that appears to regulate tuber development in potato via StBEL5-POTH1 heterodimer (Rosin *et al*., 2003). Finally, QTL underlying IHN-related traits were also found in the same LG 5 hotspot, where a homeobox-associated leucine zipper, *StHOX40* (PGSC0003DMG400030494, ST4.03ch05:4264626..4267698), was identified. This gene was found to be either up or downregulated under heat and drought stresses in potato, respectively (Li *et al*., 2019), which are believed to be two important triggers of IHN (Yencho *et al*., 2008).

Other genes, such as *StERF6* (PGSC0003DMG400016651, ST4.03ch06:33827698..33832740) and *StNAC046* (PGSC0003DMG400031266, ST4.03ch05:4946232..4948779), were characterized as transcription factors from AP2/ERF and NAC families, respectively, which have been implicated in a wide range of regulation processes in plants (Olsen *et al*., 2005; Mizoi *et al*., 2012). A gene from the NAC family, *StNAC032* (PGSC0003DMG400002824, ST4.03ch04:245125..246607), was retrieved from the LG 4 region of a major QTL for skin texture. Several genes from the NAC family, including *StNAC032*, were found to be expressed in the tuber skin and appeared to be involved in suberin and associated wax biosynthesis (Soler *et al*., 2020), which could contribute to skin texture, as well as be involved with response to drought and other biotic and abiotic stresses (Singh *et al*., 2013).

In plants, a single QTL region may be found to underlie the variation of more than one phenotype at once, which can potentially explain the correlation among traits. A QTL hotspot has been identified in the short arm of chromosome 5 of potato (Bradshaw *et al*., 2004; Danan *et al*., 2011; Li *et al*., 2018), and our B2721 population stuck to the rule. A major QTL for maturity was found to overlap loci underlying the variation for other traits such as yield, specific gravity and IHN-related resistance, as described previously (McCord *et al*., 2011a, 2011b; Schumann *et al*., 2017) and corroborated with newly analyzed phenotypic data. Pleiotropy is one possible explanation of physiologically correlated traits, such as maturity, yield and specific gravity (as a proxy of solid content, thus contributing to yield). Transcription factors are known to have this property in which a single molecular function may be involved in multiple biological processes (He and Zhang, 2006). For instance, a DNA binding element from the HAP complex, *Ghd8*, was found to play pleiotropic roles in regulating yield, flowering and plant height in rice (Yan *et al*., 2011). On the other hand, the presence of QTL for yield-related traits and pathogenic (e.g. late blight) and non-pathogenic (e.g. IHN) potato disorders in the same hotspot on LG 5 could well be due to closely linked genes. Another example from rice has suggested the multigenic nature of a major QTL for grain yield under drought, where the central role of a NAM transcription factor, *OsNAM12*.*1*, in concert with extra co-localized genes, influenced other traits such as root and panicle branching, and transpiration efficiency (Dixit *et al*., 2015).

## 5 Final Considerations

‘Atlantic’ has been widely adopted as a chipping variety, mostly due to its high specific gravity and yield, and in spite of its medium-late maturity and susceptibility to internal heat necrosis (Webb *et al*., 1978). The genetic basis of the correlation between these traits was dissected here by means of multiple QTL models in combination with a dense, fully phased genetic map. We found that not only a major QTL on LG 5 underlies the variation of these traits concomitantly, but also that the additive effects attributed to each parental haplotype appear in such a way where, in general, when an allele increases SG and FM measurements (thus towards early maturity), it decreases PY and IHN resistance (hence towards low NS and high NI), and vice-versa. *StCDF1* has been pointed out as playing a central role in this major QTL for maturity, with its day-length signaling function that regulates the tuberization pathway (Kloosterman *et al*., 2013). In addition to low rainfall in the early season and high temperatures, longer days to harvest are known to increase IHN incidence (reviewed by Yencho *et al*., 2008). Similarly, higher number of rain events seemed to contribute to the occurrence of IHN, because rainfall interferes, among other variables, with light intensity (Sterrett *et al*., 1991). Therefore, due to its circadian role, we believe that *StCDF1* indirectly participates in IHN progression.

Our list of candidate genes can be used in further dissection of QTL regions. Ideally, factors other than *StCDF1* could be targeted in benefit of yield and specific gravity increase at the expense of late maturity. One way of doing so is to focus on linked genes (e.g. *POTH1* for yield) or other non-linked QTL (e.g. the one on LG 2 for specific gravity). Moreover, the evaluation of germplasm collections in order to assess haplotypic diversity could assist the development and implementation of markers to monitor loci of interest, as previously performed for *StCDF1* in chromosome 5 (Gebhardt *et al*., 2004; Hardigan *et al*., 2017). Marker-assisted breeding for quantitative, more complex traits could be implemented upon validation of these QTL neighboring markers in breeding populations of tetraploid potato. DNA markers also can be employed in genome-wide prediction in this crop (Sverrisdóttir *et al*., 2017; Enciso-Rodriguez *et al*., 2018). In this case, both meiotic and trait genetic architecture assessments derived from our analyses can be particularly useful in refining simulations of potato and other autopolyploid crop breeding programs integrating genomics-assisted breeding (Gaynor *et al*., 2020).

## 6 Author Contributions

GCY designed the experiments and supervised the project. MJS and MEC performed field experiments and collected data. GSP, MM and ZBZ analyzed data. GSP drafted the manuscript with major contributions from MM. All authors read and approved the manuscript.

## 8 Acknowledgments

We would like to thank all the staff at the North Carolina Department of Agriculture and Consumer Services Tidewater Research Station, Plymouth, NC, for their hard work and dedication to quality research in aiding with the field aspect of this study. We also appreciate Michigan State University Potato Breeding and Genetics Program for their assistance with the Illumina Infinium^®^ 8,303 Potato Array.

## 10 Data Archiving

The datasets analyzed and scripts used in this study can be found in the supplementary material and at the study GitHub page (https://github.com/mmollina/B2721_map). Both MAPpoly (https://github.com/mmollina/MAPpoly) and QTLpoly (https://github.com/guilherme-pereira/QTLpoly) R packages are available at GitHub.

## References

Bennett MD (2004). Perspectives on polyploidy in plants – Ancient and neo. Biol J Linn Soc 82: 411–423.

Bethke PC, Nassar AMK, Kubow S, Leclerc YN, Li XQ, Haroon M, et al. (2014). History and Origin of Russet Burbank (Netted Gem) a Sport of Burbank. Am J Potato Res 91: 594–609.

Bourke PM, van Geest G, Voorrips RE, Jansen J, Kranenburg T, Shahin A, et al. (2018). polymapR—linkage analysis and genetic map construction from F1 populations of outcrossing polyploids. Bioinformatics: 1–7.

Bourke PM, Voorrips RE, Visser RGF, Maliepaard C (2015). The double-reduction landscape in tetraploid potato as revealed by a high-density linkage map. Genetics 201: 853–863.

Bradshaw JE, Hackett CA, Pande B, Waugh R, Bryan GJ (2008). QTL mapping of yield, agronomic and quality traits in tetraploid potato (Solanum tuberosum subsp. tuberosum). Theor Appl Genet 116: 193–211.

Bradshaw JE, Pande B, Bryan GJ, Hackett CA, McLean K, Stewart HE, et al. (2004). Interval mapping of quantitative trait loci for resistance to late blight [Phytophthora infestans (Mont.) de bary], height and maturity in a tetraploid population of potato (Solanum tuberosum subsp. tuberosum). Genetics 168: 983–995.

Butler DG, Cullis BR, Gilmour AR, Gogel BJ, Thompson R (2018). ASReml-R Reference Manual Version 4. Hemel Hempstead.

Chen J, Zhang F, Wang L, Leach L, Luo Z (2018). Orthogonal contrast based models for quantitative genetic analysis in autotetraploid species. New Phytol 220: 332–346.

Choudhary A, Wright L, Ponce O, Chen J, Prashar A, Sanchez-Moran E, et al. (2020). Varietal variation and chromosome behaviour during meiosis in Solanum tuberosum. Heredity (Edinb).

Churchill GA, Doerge RW (1994). Empirical threshold values for quantitative trait mapping. Genetics 138: 963–71.

Comai L (2005). The advantages and disadvantages of being polyploid. Nat Rev Genet 6: 836–846.

Danan S, Veyrieras JB, Lefebvre V (2011). Construction of a potato consensus map and QTL meta-analysis offer new insights into the genetic architecture of late blight resistance and plant maturity traits. BMC Plant Biol 11.

Dixit S, Kumar Biswal A, Min A, Henry A, Oane RH, Raorane ML, et al. (2015). Action of multiple intra-QTL genes concerted around a co-localized transcription factor underpins a large effect QTL. Sci Rep 5: 1–16.

Dumble S, Bernal EF, Villardon PG (2017). GGEBiplots: GGE Biplots with ‘ggplot2’.

Enciso-Rodriguez F, Douches D, Lopez-Cruz M, Coombs J, de los Campos G (2018). Genomic Selection for Late Blight and Common Scab Resistance in Tetraploid Potato (Solanum tuberosum). G3 Genes|Genomes|Genetics 8: 2471–2481.

Endelman JB, Carley CAS, Bethke PC, Coombs JJ, Clough ME, Silva WL, et al. (2018). Genetic Variance Partitioning and Genome-Wide Prediction with Allele Dosage Information in Autotetraploid Potato. Genetics 209: 77–87.

FAO (2020). FAOSTAT Crops.

Felcher KJ, Coombs JJ, Massa AN, Hansey CN, Hamilton JP, Veilleux RE, et al. (2012). Integration of two diploid potato linkage maps with the potato genome sequence. PLoS One 7: e36347.

Gaynor RC, Gorjanc G, Hickey JM (2020). AlphaSimR : Simulations An R-package for Breeding Program. bioRxiv: 1–21.

Gebhardt C, Ballvora A, Walkemeier B, Oberhagemann P, Schüler K (2004). Assessing genetic potential in germplasm collections of crop plants by marker-trait association: A case study for potatoes with quantitative variation of resistance to late blight and maturity type. Mol Breed 13: 93–102.

Ghislain M, Douches DS (2020). The Genes and Genomes of the Potato. In: Campos H, Ortiz O (eds) The Potato Crop, Springer, pp 139–162.

Goodstein DM, Shu S, Howson R, Neupane R, Hayes RD, Fazo J, et al. (2012). Phytozome: a comparative platform for green plant genomics. Nucleic Acids Res 40: D1178--D1186.

Hackett CA, Boskamp B, Vogogias A, Preedy KF, Milne I, Vogogias T, et al. (2016). TetraploidSNPMap: software for linkage analysis and QTL mapping in autotetraploid populations using SNP dosage data. J Hered 108: 438–442.

Hackett CA, Bradshaw JE, Bryan GJ (2014). QTL mapping in autotetraploids using SNP dosage information. Theor Appl Genet 127: 1885–1904.

Hämäläinen JH, Watanabe KN, Valkonen JPT, Arihara A, Plaisted RL, Pehu E, et al. (1997). Mapping and marker-assisted selection for a gene for extreme resistance to potato virus Y. Theor Appl Genet 94: 192–197.

Hardigan MA, Laimbeer FPE, Newton L, Crisovan E, Hamilton JP, Vaillancourt B, et al. (2017). Genome diversity of tuber-bearing Solanum uncovers complex evolutionary history and targets of domestication in the cultivated potato. Proc Natl Acad Sci U S A 114: E9999–E10008.

He X, Zhang J (2006). Toward a molecular understanding of pleiotropy. Genetics 173: 1885–1891.

Holland JB, Nyquist WE, Cervantes-Martínez CT (2003). Estimating and interpreting heritability for plant breeding: an update. In: Janick J (ed) Plant Breeding Reviews, John Wiley & Sons: New Jersey Vol 22, pp 9–112.

Huang K, Wang T, Dunn DW, Zhang P, Cao X, Liu R, et al. (2019). Genotypic Frequencies at Equilibrium for Polysomic Inheritance Under Double-Reduction. G3 Genes|Genomes|Genetics 9: 1693–1706.

Jong H (1981). Inheritance of russeting in cultivated diploid potatoes. Potato Res 24: 309–313.

Kempthorne O (1955). The correlation between relatives in a simple autotetraploid population. Genetics 40: 168–174.

Kloosterman B, Abelenda JA, Gomez MDMC, Oortwijn M, De Boer JM, Kowitwanich K, et al. (2013). Naturally occurring allele diversity allows potato cultivation in northern latitudes. Nature 495: 246–250.

Li W, Dong J, Cao M, Gao X, Wang D, Liu B, et al. (2019). Genome-wide identification and characterization of HD-ZIP genes in potato. Gene 697: 103–117.

Li X, Xu J, Duan S, Zhang J, Bian C, Hu J, et al. (2018). Mapping and QTL analysis of early-maturity traits in tetraploid potato (Solanum tuberosum L.). Int J Mol Sci 19.

Luo ZW, Zhang RM, Kearsey MJ (2004). Theoretical basis for genetic linkage analysis in autotetraploid species. PNAS 101: 7040–7045.

Mann H, Iorizzo M, Gao L, D’Agostino N, Carputo D, Chiusano ML, et al. (2011). Molecular Linkage Maps: Strategies, Resources and Achievements. In: Bradeen JM, Kole C (eds) Genetics, Genomics and Breeding of Potato, CRC Press: Enfield, pp 68–89.

Manrique-Carpintero NC, Coombs JJ, Cui Y, Veilleux RE, Robin Buell C, Douches D (2015). Genetic map and QTL analysis of agronomic traits in a diploid potato population using single nucleotide polymorphism markers. Crop Sci 55: 2566–2579.

Massa AN, Manrique-Carpintero NC, Coombs J, Haynes KG, Bethke PC, Brandt TL, et al. (2018). Linkage analysis and QTL mapping in a tetraploid russet mapping population of potato. BMC Genet 19: 1–13.

Massa AN, Manrique-Carpintero NC, Coombs JJ, Zarka DG, Boone AE, Kirk WW, et al. (2015). Genetic linkage mapping of economically important traits in cultivated tetraploid potato (Solanum tuberosum L.). G3 Genes, Genomes, Genet 5: 2357–2364.

McCord PH, Sosinski BR, Haynes KG, Clough ME, Yencho GC (2011a). QTL mapping of internal heat necrosis in tetraploid potato. Theor Appl Genet 122: 129–142.

McCord PH, Sosinski BR, Haynes KG, Clough ME, Yencho GC (2011b). Linkage mapping and QTL analysis of agronomic traits in tetraploid potato (Solanum tuberosum subsp. tuberosum). Crop Sci 51: 771–785.

Mengist MF, Alves S, Griffin D, Creedon J, McLaughlin MJ, Jones PW, et al. (2018). Genetic mapping of quantitative trait loci for tuber-cadmium and zinc concentration in potato reveals associations with maturity and both overlapping and independent components of genetic control. Theor Appl Genet 131: 929–945.

Mizoi J, Shinozaki K, Yamaguchi-Shinozaki K (2012). AP2/ERF family transcription factors in plant abiotic stress responses. Biochim Biophys Acta – Gene Regul Mech 1819: 86–96.

Mollinari M, Garcia AAF (2019). Linkage Analysis and Haplotype Phasing in Experimental Autopolyploid Populations with High Ploidy Level Using Hidden Markov Models. G3 Genes|Genomes|Genetics 9: 3297–3314.

Mollinari M, Olukolu BA, Pereira GS, Khan A, Gemenet D, Yencho GC, et al. (2020). Unraveling the Hexaploid Sweetpotato Inheritance Using Ultra-Dense Multilocus Mapping. G3 Genes|Genomes|Genetics 10: 281–292.

Olsen AN, Ernst HA, Leggio L Lo, Skriver K (2005). NAC transcription factors: structurally distinct, functionally diverse. Trends Plant Sci 10: 79–87.

Pereira G da S, Gemenet DC, Mollinari M, Olukolu BA, Wood JC, Diaz F, et al. (2020). Multiple QTL Mapping in Autopolyploids: A Random-Effect Model Approach with Application in a Hexaploid Sweetpotato Full-Sib Population. Genetics 215: 579–595.

Preedy KF, Hackett CA (2016). A rapid marker ordering approach for high-density genetic linkage maps in experimental autotetraploid populations using multidimensional scaling. Theor Appl Genet 129: 2117–2132.

Qu L, Guennel T, Marshall SL (2013). Linear score tests for variance components in linear mixed models and applications to genetic association studies. Biometrics 69: 883–892.

R Core Team (2020). R: A Language and Environment for Statistical Computing.

Rak K, Bethke PC, Palta JP (2017). QTL mapping of potato chip color and tuber traits within an autotetraploid family. Mol Breed 37.

Ramakrishnan AP, Ritland CE, Blas Sevillano RH, Riseman A (2015). Review of Potato Molecular Markers to Enhance Trait Selection. Am J Potato Res 92: 455–472.

Revelle W (2018). psych: Procedures for Psychological, Psychometric, and Personality Research.

Rosin FM, Hart JK, Horner HT, Davies PJ, Hannapel DJ (2003). Overexpression of a Knotted-Like Homeobox Gene of Gibberellin Accumulation 1. Plant Physiol 132: 106–117.

Ruiz E, Jackson S, Cimentada J (2019). corrr: Correlations in R.

Schmitz Carley CA, Coombs JJ, Douches DS, Bethke PC, Palta JP, Novy RG, et al. (2017). Automated tetraploid genotype calling by hierarchical clustering. Theor Appl Genet 130: 717–726.

Schultz L, Cogan NOI, Mclean K, Dale MFB, Bryan GJ, Forster JW, et al. (2012). Evaluation and implementation of a potential diagnostic molecular marker for H1-conferred potato cyst nematode resistance in potato (Solanum tuberosum L.). Plant Breed 131: 315–321.

Schumann MJ, Zeng ZB, Clough ME, Yencho GC (2017). Linkage map construction and QTL analysis for internal heat necrosis in autotetraploid potato. Theor Appl Genet 130: 2045–2056.

Shaner G, Finney RE (1977). The Effect of Nitrogen Fertilization on the Expression of Slow – Mildewing Resistance in Knox Wheat. Phytopathology 67: 1051–1056.

Sharma SK, Bolser D, de Boer J, Sonderkaer M, Amoros W, Carboni MF, et al. (2013). Construction of reference chromosome-scale pseudomolecules for potato: integrating the potato genome with genetic and physical maps. G3 Genes|Genomes|Genetics 3: 2031–47.

Singh AK, Sharma V, Pal AK, Acharya V, Ahuja PS (2013). Genome-wide organization and expression profiling of the NAC transcription factor family in potato (solanum tuberosum L.). DNA Res 20: 403–423.

Slater AT, Cogan NOI, Hayes BJ, Schultz L, Dale MFB, Bryan GJ, et al. (2014). Improving breeding efficiency in potato using molecular and quantitative genetics. Theor Appl Genet 127: 2279–2292.

Soler M, Verdaguer R, Fernández-Piñán S, Company-Arumí D, Boher P, Góngora-Castillo E, et al. (2020). Silencing against the conserved NAC domain of the potato StNAC103 reveals new NAC candidates to repress the suberin associated waxes in phellem. Plant Sci 291: 110360.

Sterrett SB, Lee GS, Henninger MR, Lentner M (1991). Predictive Model for Onset and Development of Internal Heat Necrosis of ‘Atlantic’ Potato. J Am Soc Hortic Sci 116: 701–705.

Sverrisdóttir E, Byrne S, Sundmark EHR, Johnsen Hø, Kirk HG, Asp T, et al. (2017). Genomic prediction of starch content and chipping quality in tetraploid potato using genotyping-by-sequencing. Theor Appl Genet 130: 2091–2108.

Webb RE, Wilson DR, Shumaker JR, Graves B, Henninger MR, Watts J, et al. (1978). Atlantic: A new potato variety with high solids, good processing quality, and resistance to pests. Am Potato J 55: 141–145.

Wei T, Simko V, Levy M, Xie Y, Jin Y, Zemla J (2017). corrplot: Visualization of a Correlation Matrix.

Wickham H (2016). ggplot2: elegant graphics for data analysis. Springer.

Xu X, Pan S, Cheng S, Zhang B, Mu D, Ni P, et al. (2011). Genome sequence and analysis of the tuber crop potato. Nature 475: 189–195.

Yan W, Kang MS (2003). GGE Biplot Analysis: a graphical tool for breeders, geneticists, and agronomists. CRC Press: Boca Raton.

Yan WH, Wang P, Chen HX, Zhou HJ, Li QP, Wang CR, et al. (2011). A major QTL, Ghd8, plays pleiotropic roles in regulating grain productivity, plant height, and heading date in rice. Mol Plant 4: 319–330.

Ye J, Zhang Y, Cui H, Liu J, Wu Y, Cheng Y, et al. (2018). WEGO 2.0: A web tool for analyzing and plotting GO annotations, 2018 update. Nucleic Acids Res 46: W71–W75.

Yencho GC, McCord PH, Haynes KG, Sterrett SBR (2008). Internal heat necrosis of potato – A review. Am J Potato Res 85: 69–76.

Zielinski M-L, Scheid OM (2012). Meiosis in Polyploid Plants. In:Soltis PS, Soltis DE (eds) Polyploidy and Genome Evolution, Springer: Berlin, Heidelberg, pp 33–55.

Zou F, Fine JP, Hu J, Lin DY (2004). An efficient resampling method for assessing genome-wide statistical significance in mapping quantitative trait loci. Genetics 168: 2307–2316.

Zych K, Gort G, Maliepaard CA, Jansen RC, Voorrips RE (2019). FitTetra 2.0 – Improved genotype calling for tetraploids with multiple population and parental data support. BMC Bioinformatics 20: 1–8.

