## Supporting Information for "The recombination landscape and multiple QTL mapping in a *Solanum tuberosum* cv. ‘Atlantic’-derived F_1_ population"

Article acceptance date: Click here to enter a date.

The following Supporting Information is available for this article:

**Fig. S1** Correlation between adjusted means for B2721 traits.

**Fig. S2** Overview of SNP filtered data for B2721 map construction.

**Fig. S3** B2721 genetic map representation.

**Fig. S4** Genetic map and reference genome scatterplots.

**Fig. S5** Homolog pairing assessment in B2721 mapping population.

**Fig. S6** Random-effect multiple interval mapping QTL profiles for adjusted means.

**Fig. S7** Random-effect multiple interval mapping QTL profiles for maturity-corrected phenotypes.

**Fig. S8** Fixed-effect interval mapping QTL profiles for adjusted means.

**Fig. S9** Fixed-effect interval mapping QTL profiles for maturity-corrected phenotypes.

**Fig. S10** Correlation between maturity-corrected phenotypes and foliage maturity.

**Fig. S11** Correlation between QTL-based predictions.

**Fig. S12** Gene Ontology enrichment for genes within QTL regions.

**Table S1** B2721 genetic map summary.

**Table S2** Linear regression for maturity-corrected phenotypes.

**Table S3** Marker information on QTL detected via REMIM.

**Table S4** QTL information detected via FEIM.

**File S1** B2721 adjusted means and maturity-corrected phenotypes. (CSV)

**File S2** Correlation between B2721 mapping population phenotypes. (XLXS)

**File S3** B2721 normalized intensities $(x,y)$ from Illumina Infinium^®^ 8,303 Potato Array. (CSV)

**File S4** Illumina Infinium^®^ 8,303 Potato Array sequence BLAST alignment against the *Solanum tuberosum* genome v. 4.03. (CSV)

**File S5** Linkage map information for B2721 mapping population. (CSV)

**File S6** QTL additive effect effects for QTL detected via REMIM. (CSV)

**File S7** List of 533 annotated genes from *Solanum tuberosum* v. 4.03 genome related with B2721 QTL regions. (CSV)

**Fig. S1** Correlation between adjusted means for B2721 traits evaluated over four years (2006-8 and 2014) based on pairwise complete observations. Traits: plant yield (PY), foliage maturity (FM), specific gravity (SG), dry matter (DM), skin texture (ST), and internal heat necrosis severity (NS) and intensity (NI).

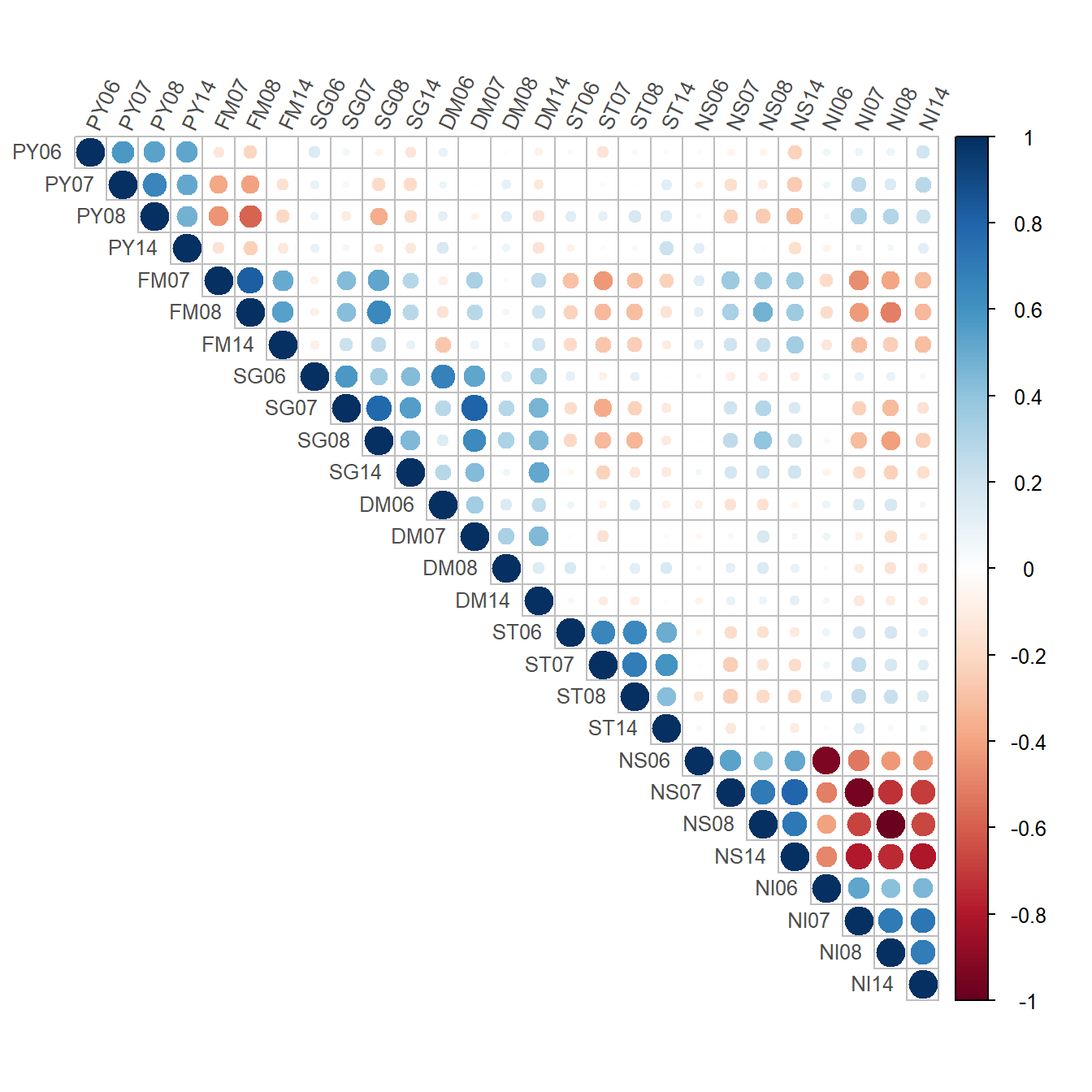

**Fig. S2** Filtered data used to build the B2721 population map: 4,812 SNPs scored in 156 individuals. The barplot indicates the dosage combinations for all markers: 1-0 indicates 0 doses for parent ‘Atlantic’ and 1 dose for parent B1829-5, and so on. The blue dots indicate the $\log_{10}(P)$ for a chi-square test of segregation distortion under Mendelian inheritance. The colored panel indicates the distribution of the dosages in the population.

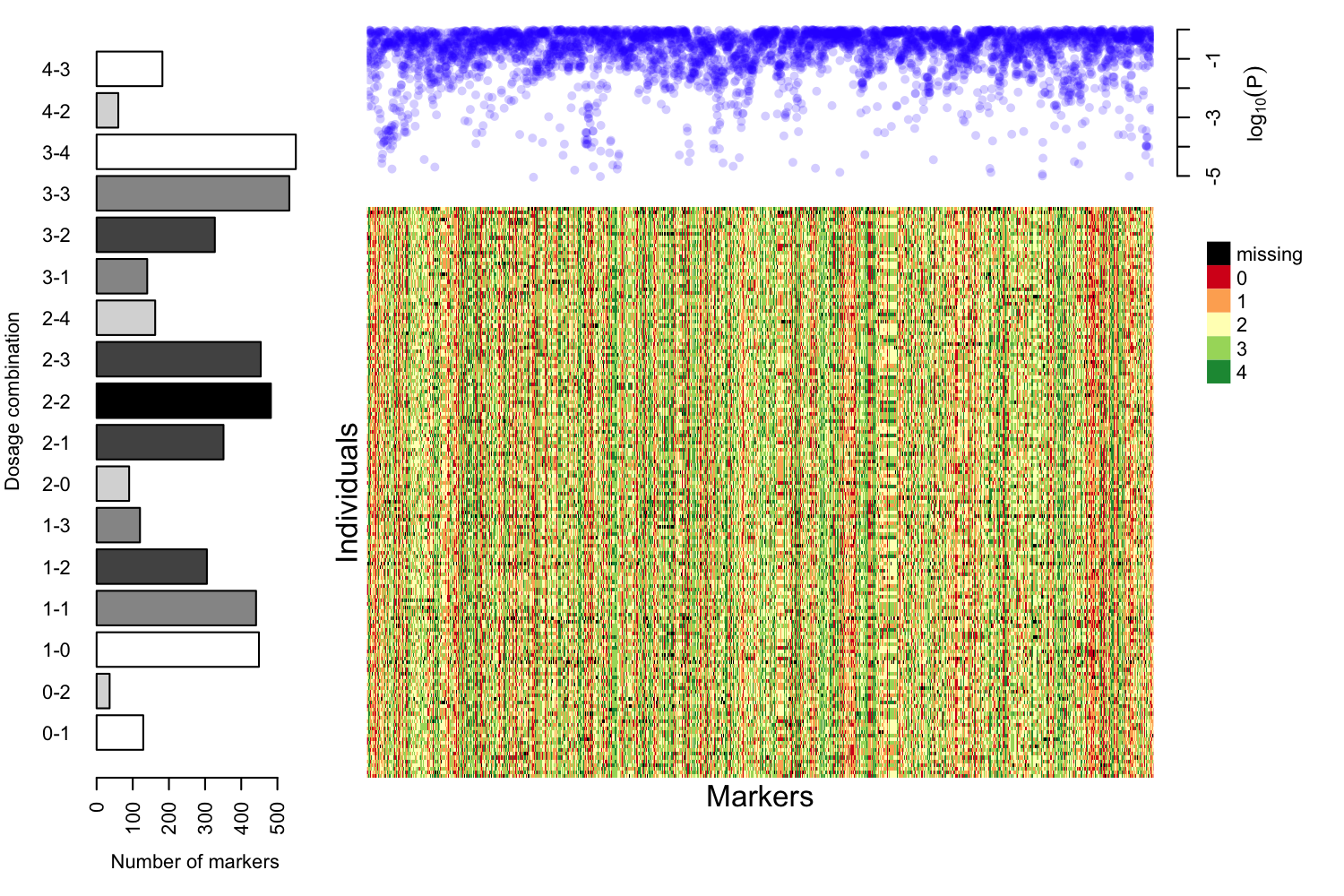

**Fig. S3** Genetic map of the B2721 population, with black vertical lines representing SNPs in their respective positions.

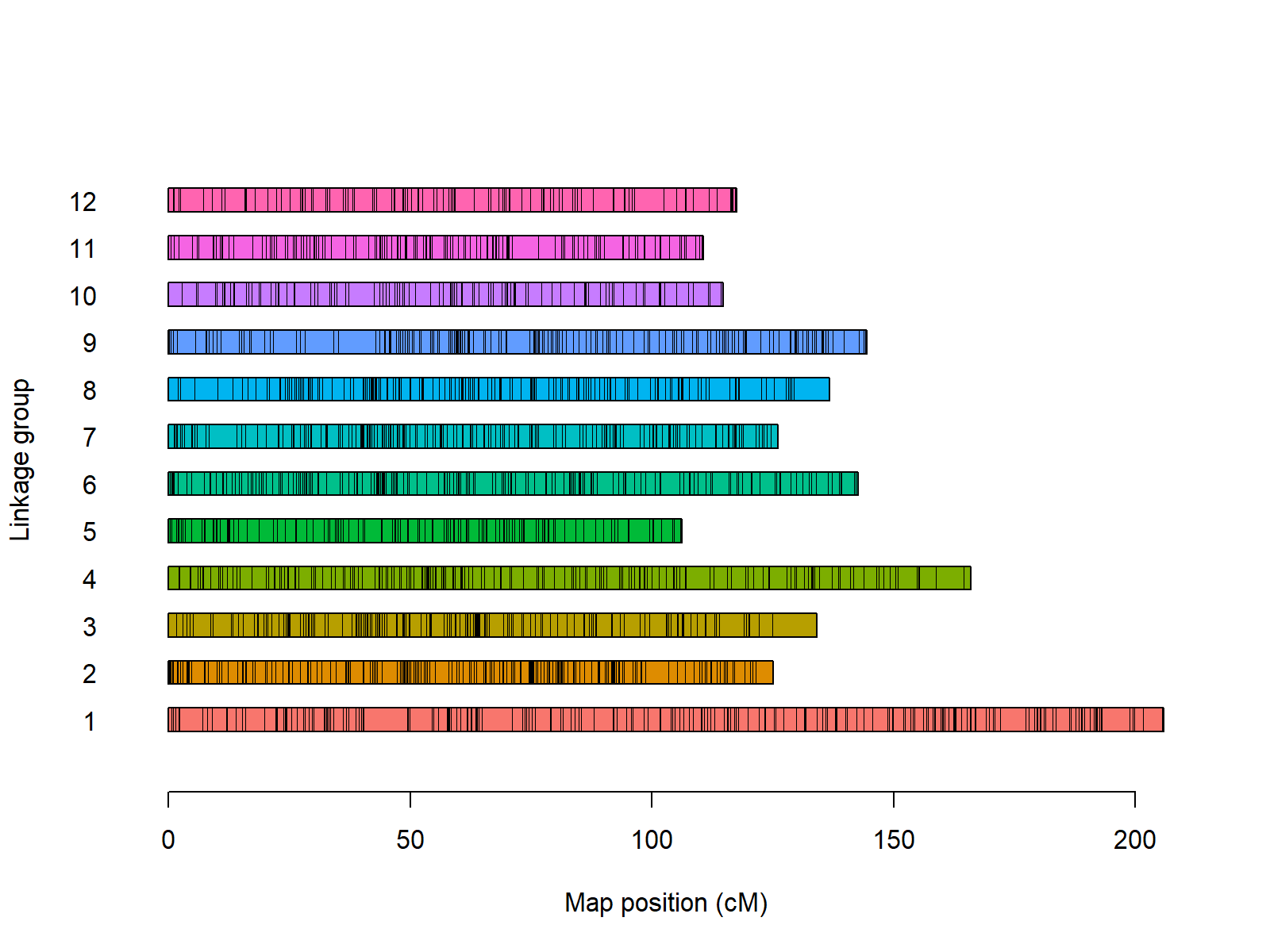

**Fig. S4** Scatterplots of the B2721 genetic map (in centiMorgans, cM) versus *Solanum tuberosum* v. 4.03 reference genome (in mega base pairs, Mbp).

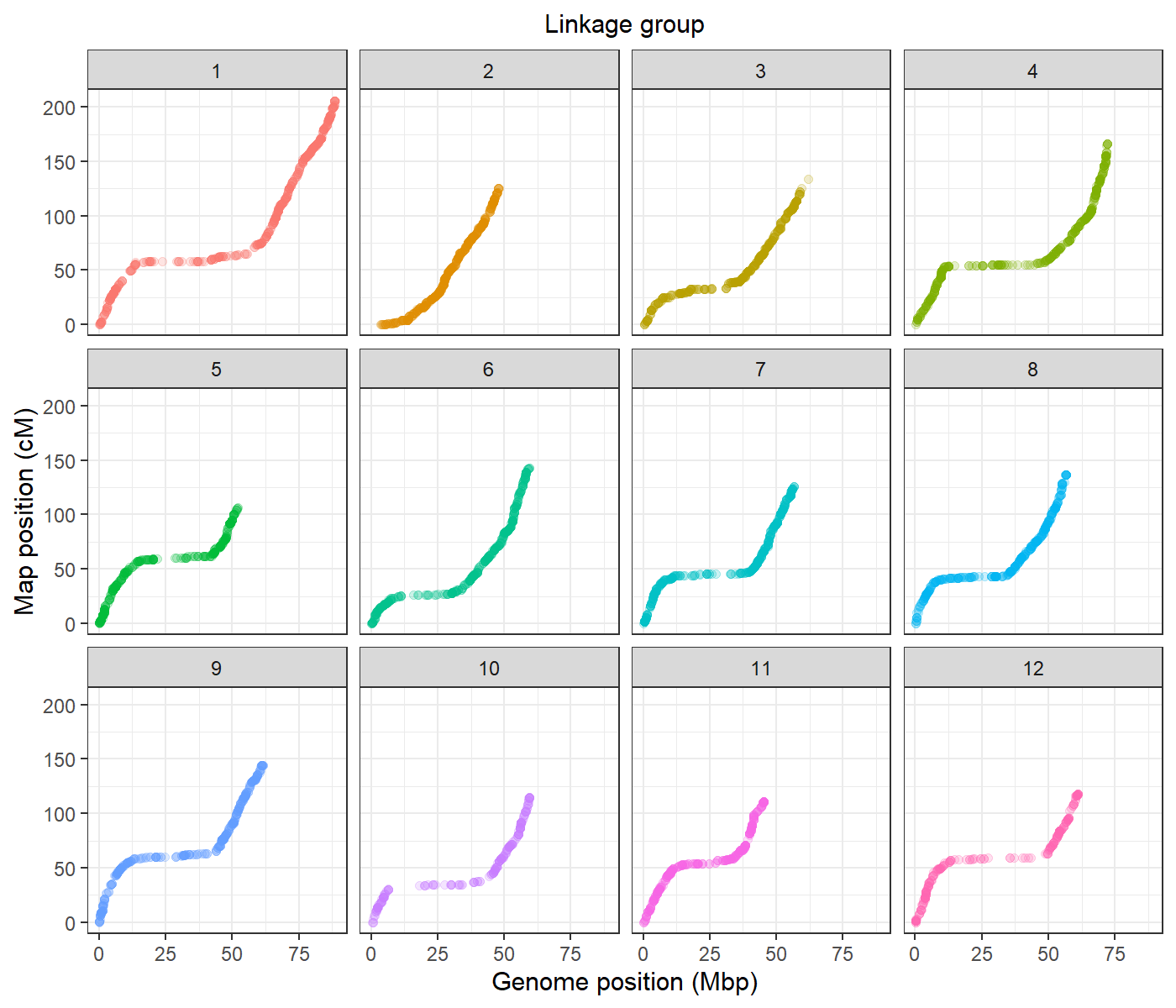

**Fig. S5** Homolog pairing assessment in B2721 mapping population. (A) Probabilistic pairing profiles, where parental homologs (‘Atlantic’ = *a* through *d*, B1829-5 = *e* through *f*) are paired according to the following notation: e.g. *ab*/*cd* where homolog *a* paired with *b*, and *c* paired with *d*. The dashed line is the expected probability under random pairing (1/3). (B) $P$-values associated to the chi-square test with the null hypothesis that all pairing configurations have the same probability. The dashed line represents $P=0.01$.

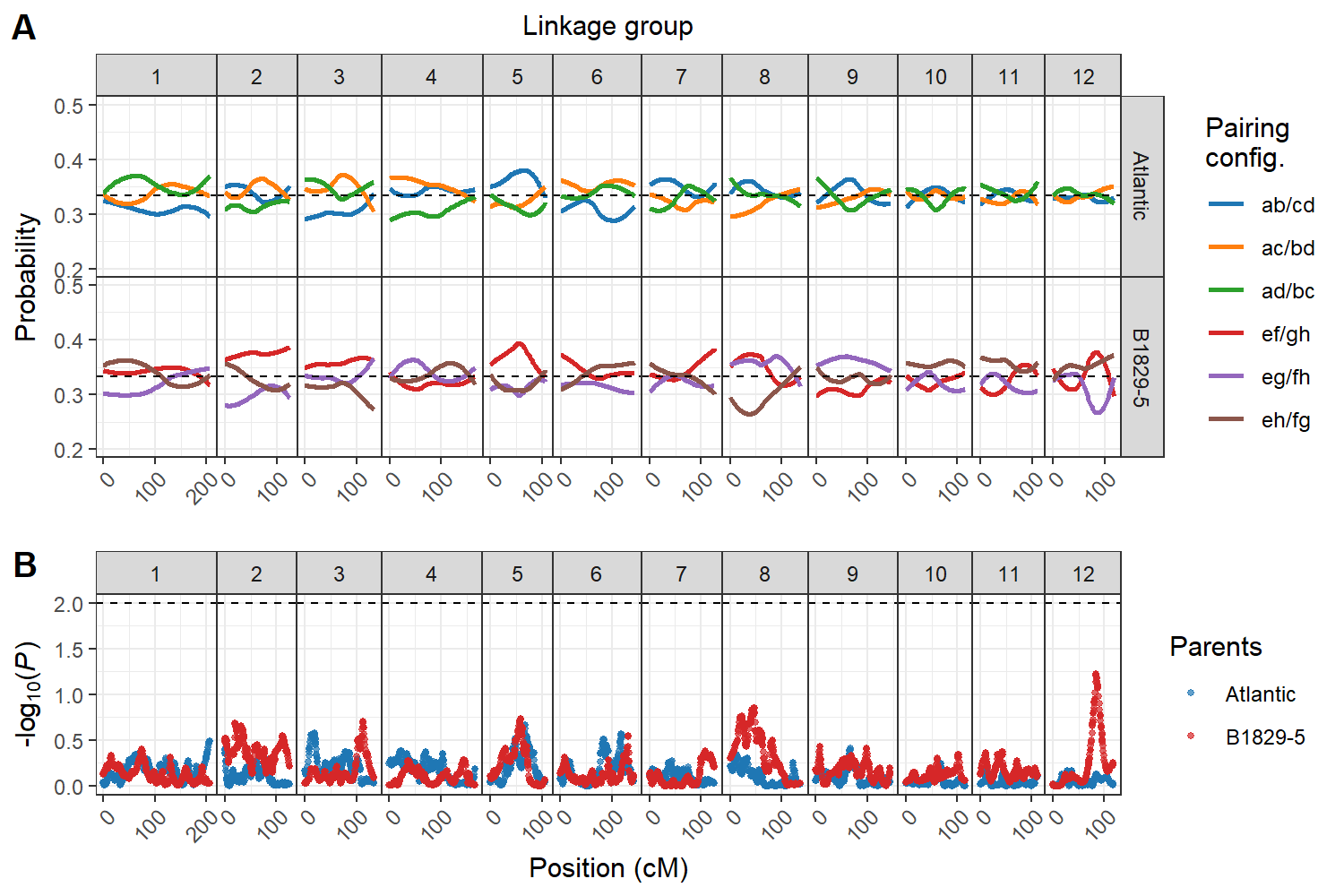

**Fig. S6** Logarithm of the $P$-values [$LOP=-\log_{10} (P)$] profiles from random-effect multiple interval mapping (REMIM) for seven B2721 traits evaluated over four years (2006-8 and 2014). QTL peak locations are represented by triangles and their ~95% support intervals by light-shaded rectangles. Traits: plant yield (PY), foliage maturity (FM), specific gravity (SG), dry matter (DM), skin texture (ST), and internal heat necrosis severity (NS) and intensity (NI).

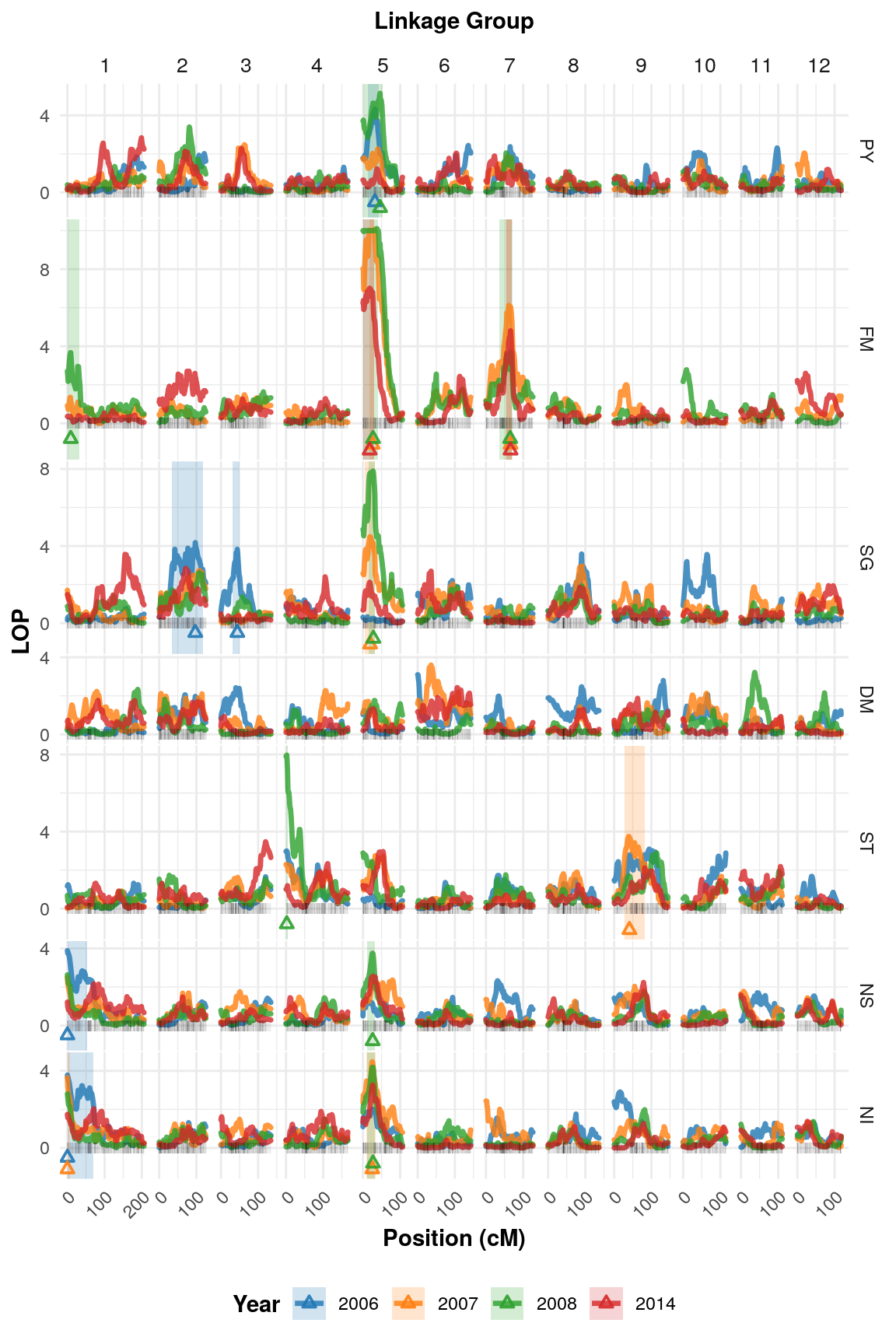

**Fig. S7** Logarithm of the $P$-values [$LOP=-\log_{10} (P)$] profiles from random-effect multiple interval mapping (REMIM) for foliage maturity (FM) corrected phenotypes evaluated over three years (2007, 2008 and 2014). QTL peak locations are represented by triangles and their ~95% support intervals by light-shaded rectangles. Traits: plant yield (PY), specific gravity (SG), dry matter (DM), skin texture (ST), and internal heat necrosis severity (NS) and intensity (NI).

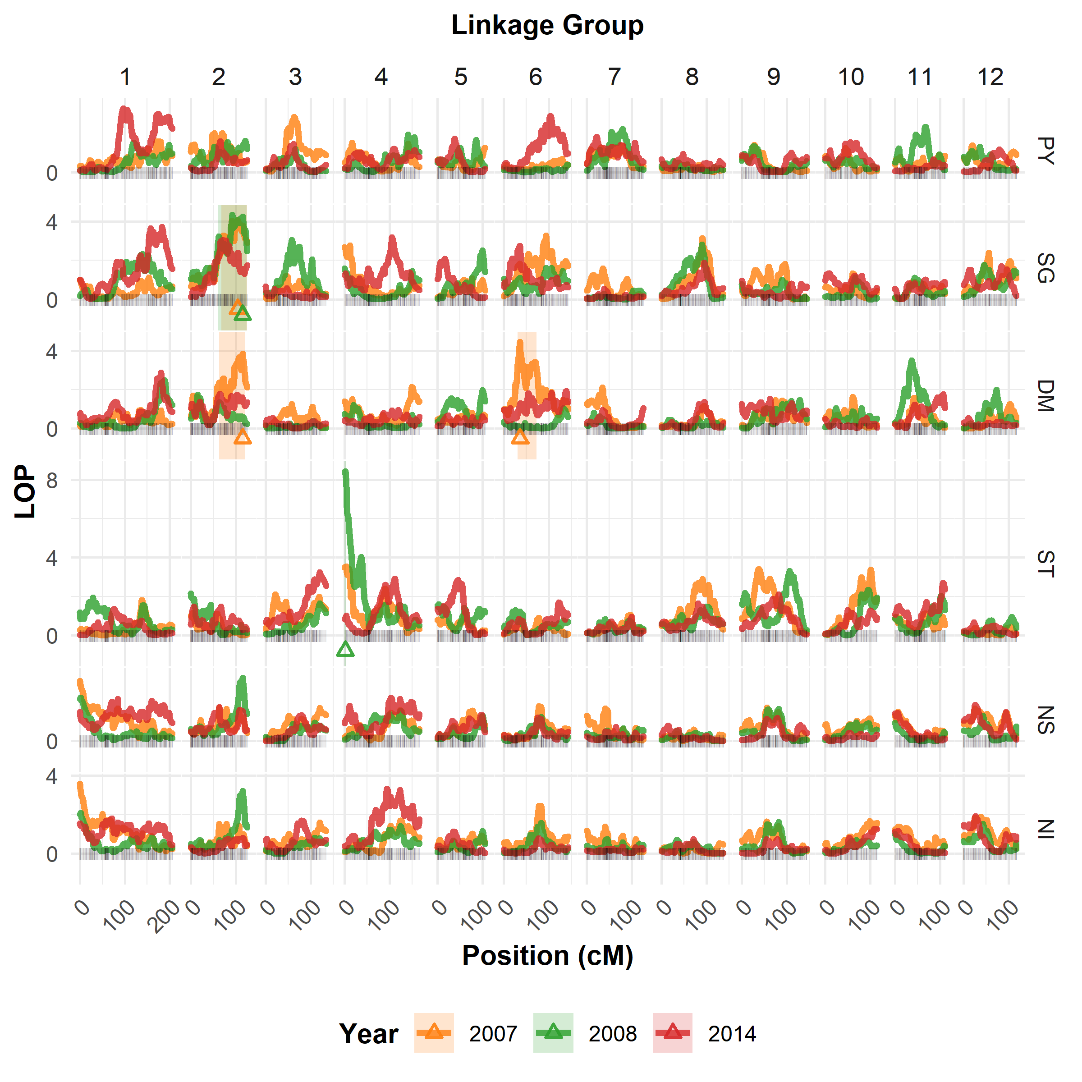

**Fig. S8** Logarithm of the odds (LOD) profiles from fixed-effect interval mapping (FEIM) for seven B2721 traits evaluated over four years (2006-8 and 2014). QTL peak locations are represented by triangles and their ~95% support intervals by light-shaded rectangles. Dashed lines denote the permutation-based thresholds. Traits: plant yield (PY), foliage maturity (FM), specific gravity (SG), dry matter (DM), skin texture (ST), and internal heat necrosis severity (NS) and intensity (NI).

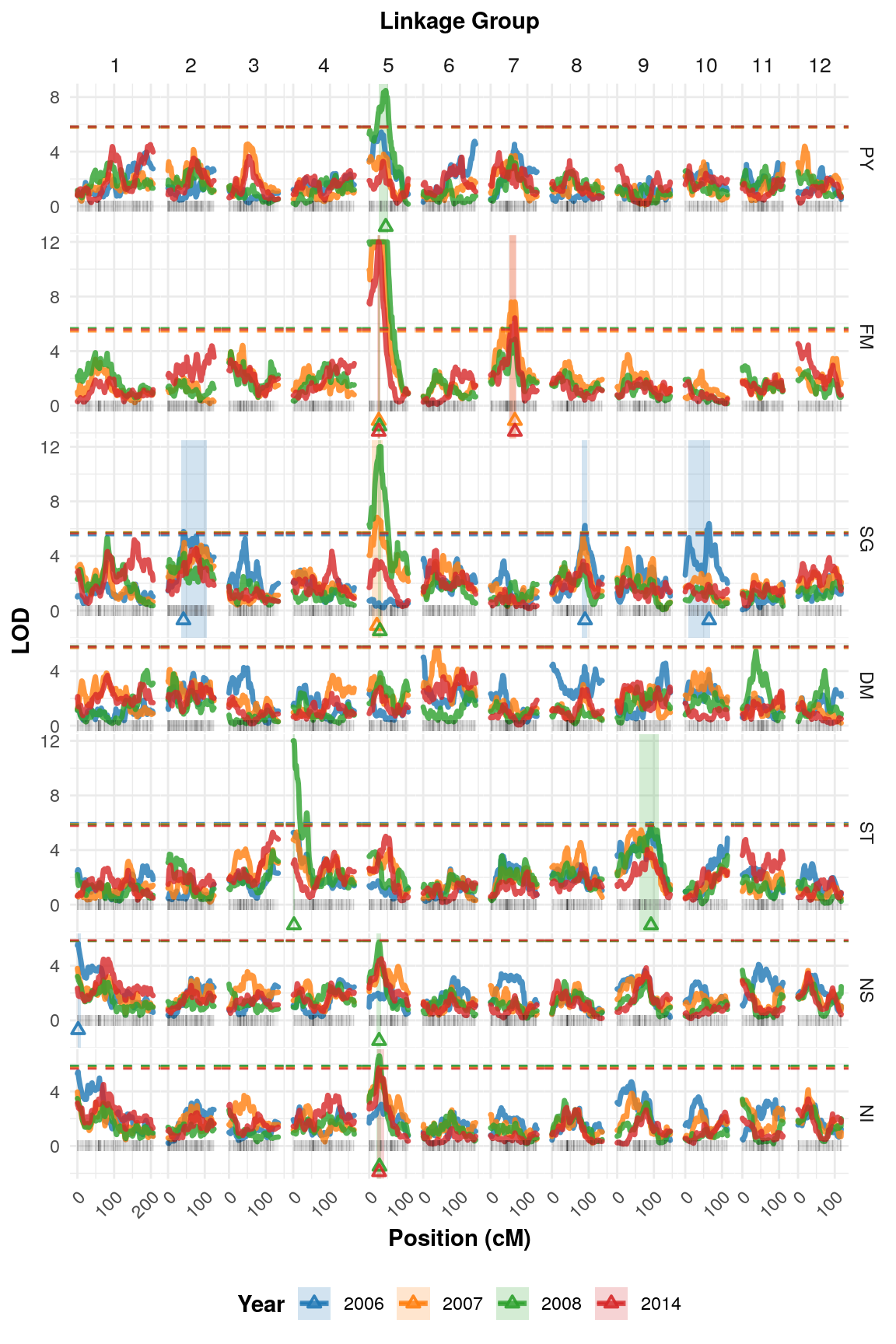

**Fig. S9** Logarithm of the odds (LOD) profiles from fixed-effect interval mapping (FEIM) for foliage maturity (FM) corrected phenotypes evaluated over three years (2007, 2008 and 2014). QTL peak locations are represented by triangles and their ~95% support intervals by light-shaded rectangles. Dashed lines denote the permutation-based thresholds. Traits: plant yield (PY), specific gravity (SG), dry matter (DM), skin texture (ST), and internal heat necrosis severity (NS) and intensity (NI).

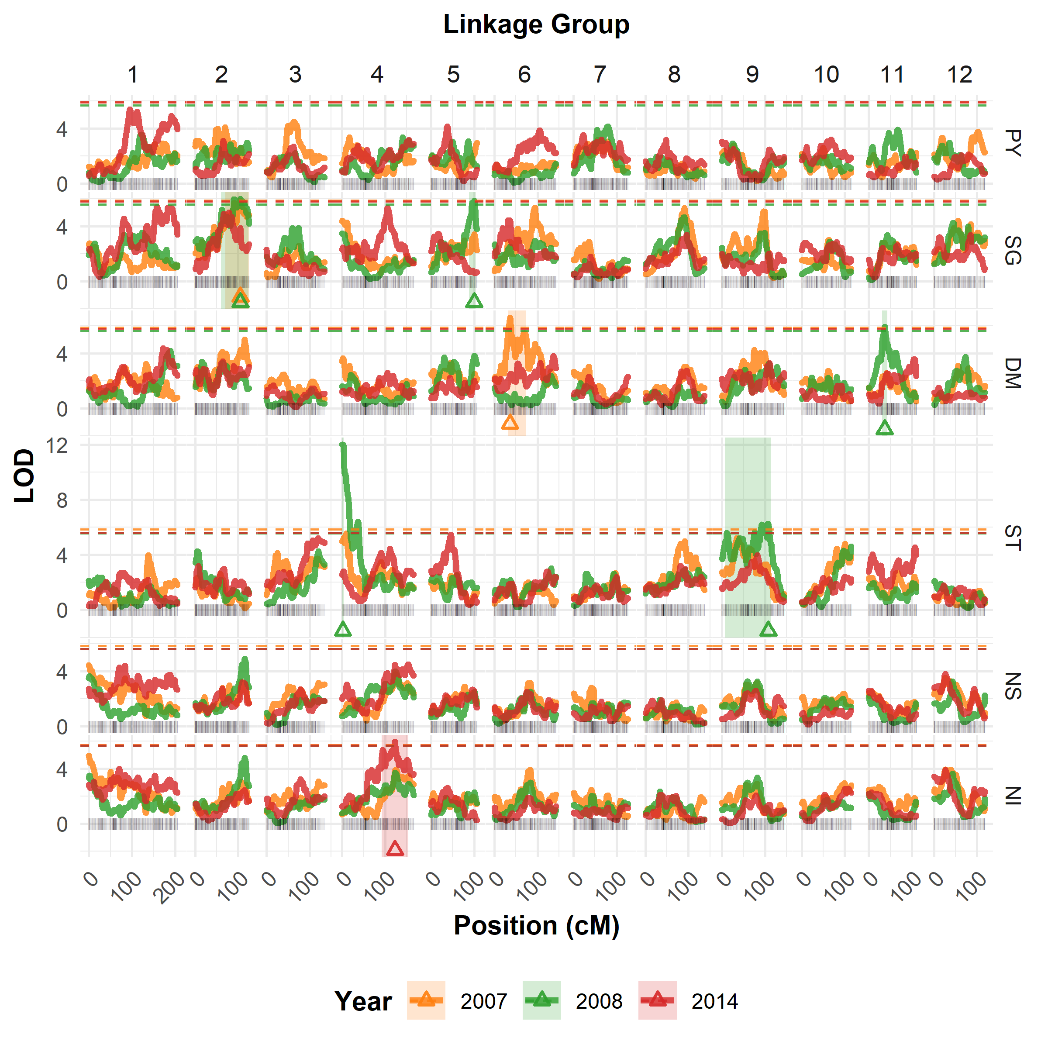

**Fig. S10** Correlations between maturity-corrected phenotypes evaluated over three years (2007, 2008 and 2014). Traits: plant yield (PY), foliage maturity (FM), specific gravity (SG), dry matter (DM), skin texture (ST), and internal heat necrosis severity (NS) and intensity (NI). * indicates maturity-corrected phenotypes.

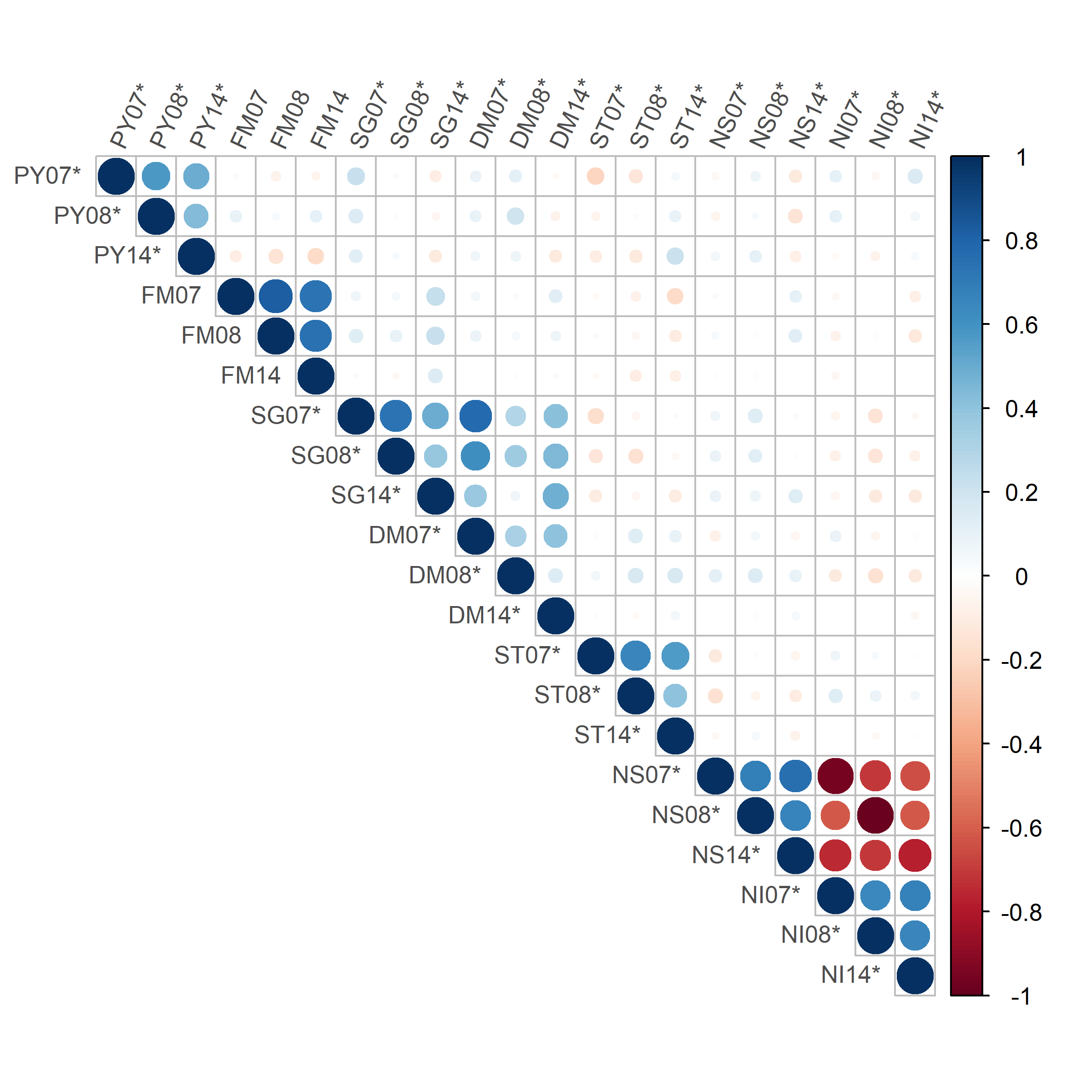

**Fig. S11** Correlations between adjusted means (upper diagonal) and QTL-based predictions (lower diagonal) for B2721 traits (only those with identified QTL) evaluated over four years (2006-8 and 2014). Traits: plant yield (PY), foliage maturity (FM), specific gravity (SG), dry matter (DM), skin texture (ST), and internal heat necrosis severity (NS) and intensity (NI). * indicates maturity-corrected phenotypes.

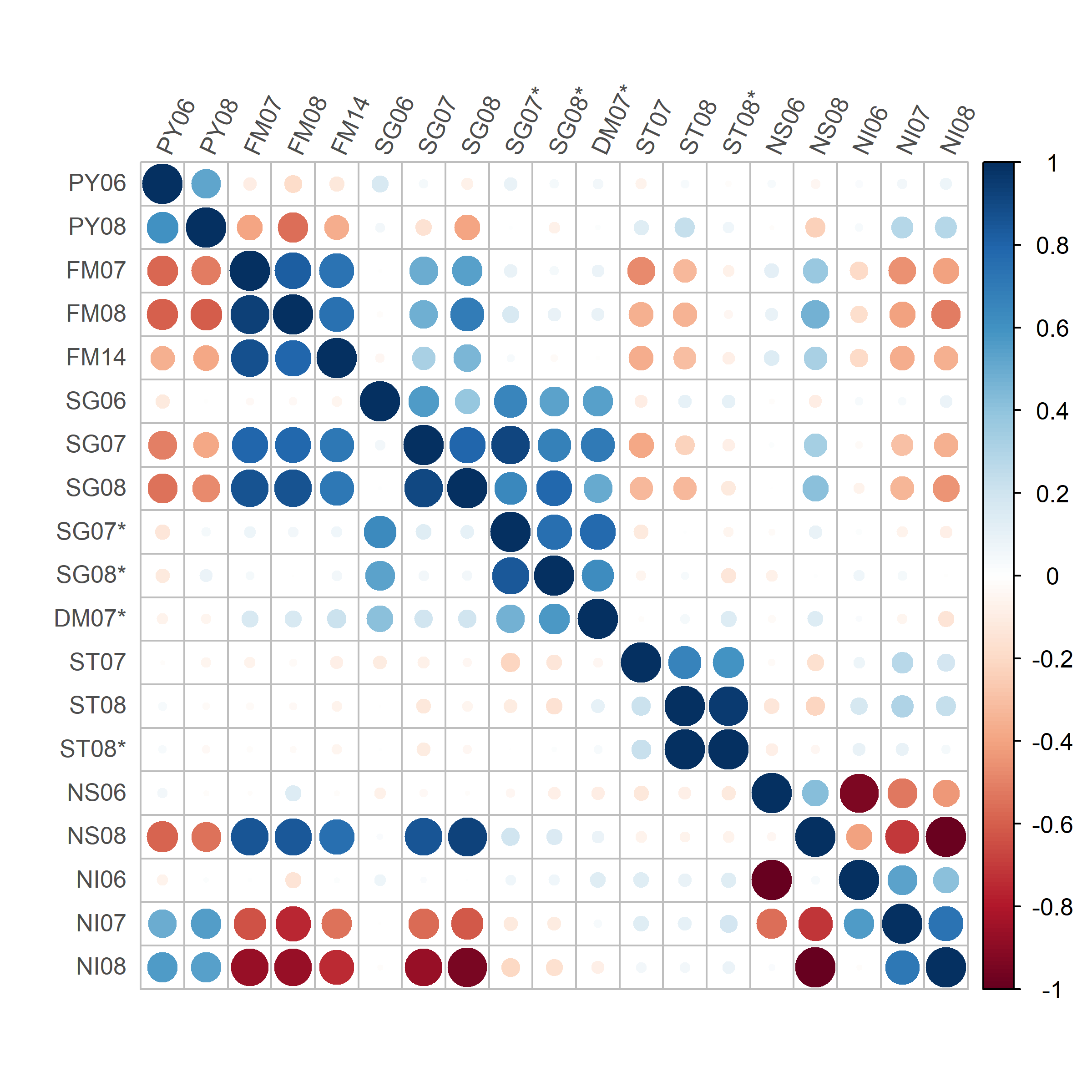

**Fig. S12** Enriched Gene Ontology (GO) terms for 533 annotated genes within our QTL regions.

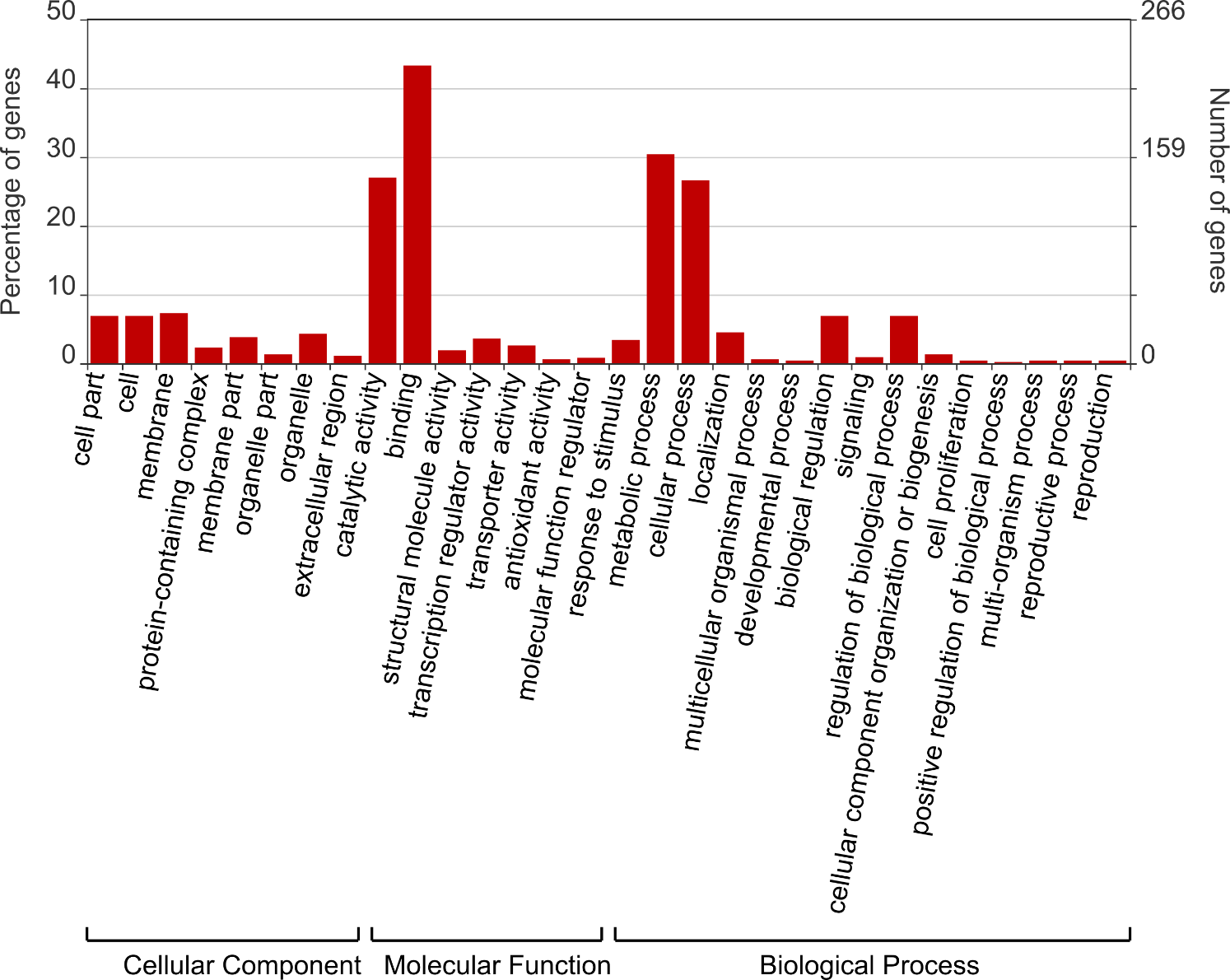

**Table S1** B2721 genetic map summary.

| Linkage  group | Length  (cM) | Number of SNPs | | | Total | SNPs/cM |
| --- | --- | --- | --- | --- | --- | --- |
|  |  | Simplex | Double-  simplex | Multiplex |  |  |
| 1 | 205.88 | 102 | 122 | 252 | 476 | 2.31 |
| 2 | 125.11 | 131 | 139 | 160 | 430 | 3.44 |
| 3 | 134.07 | 151 | 21 | 210 | 382 | 2.85 |
| 4 | 165.90 | 114 | 87 | 224 | 425 | 2.56 |
| 5 | 106.20 | 124 | 53 | 134 | 311 | 2.93 |
| 6 | 142.70 | 73 | 75 | 249 | 397 | 2.78 |
| 7 | 126.09 | 136 | 94 | 184 | 414 | 3.28 |
| 8 | 136.68 | 61 | 98 | 183 | 342 | 2.50 |
| 9 | 144.49 | 76 | 101 | 179 | 356 | 2.46 |
| 10 | 114.67 | 47 | 36 | 124 | 207 | 1.81 |
| 11 | 110.66 | 95 | 52 | 173 | 320 | 2.89 |
| 12 | 117.54 | 115 | 32 | 78 | 225 | 1.91 |
| Total | 1,629.99 | 1,225 | 910 | 2,150 | 4,285 | 2.64 |

**Table S2** Simple linear regression results of maturity-corrected phenotypes for year 2007, 2008 and 2014 where foliage maturity (FM) data was available.

|  | Trait^a^ | $\beta_{0}$ | $\beta_{1}$ | | | |
| --- | --- | --- | --- | --- | --- | --- |
|  |  |  | Estimate | Std. error | $t$ value | $\Pr(>\vert t\vert)$^b^ |
| FM07 | PY07 | 0.638 | –0.00258 | 0.00051 | –5.053 | 1.22e–06*** |
|  | SG07 | 1.065 | 0.00014 | 0.00002 | 5.951 | 1.74e–08*** |
|  | DM07 | 19.242 | 0.02013 | 0.00488 | 4.124 | 6.08e–05*** |
|  | ST07 | 2.516 | –0.01780 | 0.00293 | –6.069 | 9.60e–09*** |
|  | NS07 | 7.201 | 0.01710 | 0.00345 | 4.957 | 1.87e–06*** |
|  | NI07 | 61.588 | –0.57600 | 0.08746 | –6.586 | 6.74e–10*** |
| FM08 | PY08 | 0.430 | –0.00190 | 0.00022 | –8.782 | 2.91e–15*** |
|  | SG08 | 1.057 | 0.00017 | 0.00002 | 10.463 | 1.09e–19*** |
|  | DM08 | 19.258 | 0.00593 | 0.00801 | 0.740 | 4.61e–01 |
|  | ST08 | 2.027 | –0.00997 | 0.00250 | –3.982 | 1.05e–04*** |
|  | NS08 | 8.487 | 0.00472 | 0.00071 | 6.642 | 5.01e–10*** |
|  | NI08 | 22.504 | –0.20462 | 0.02838 | –7.211 | 2.35e–11*** |
| FM14 | PY14 | 0.549 | –0.00268 | 0.00169 | –1.584 | 1.15e–01 |
|  | SG14 | 1.059 | 0.00013 | 0.00010 | 1.232 | 2.20e–01 |
|  | DM14 | 13.871 | 0.06831 | 0.02903 | 2.353 | 1.99e–02* |
|  | ST14 | 3.136 | –0.01130 | 0.00886 | –1.276 | 2.04e–01 |
|  | NS14 | 1.220 | 0.07873 | 0.01756 | 4.484 | 1.45e–05*** |
|  | NI14 | 152.732 | –1.49138 | 0.37342 | –3.994 | 1.02e–04*** |

^a^Traits: plant yield (PY), specific gravity (SG), skin texture (ST), and internal heat necrosis severity (NS) and intensity (NI).

^b^***$P<0.001$, *$P<0.05$.

**Table S3** Illumina Infinium^®^ 8,303 Potato Array marker names (solcap_snp) and their respective *Solanum tuberosum* v. 4.03 (ST4.03) genome positions, in base pairs (bp), on the left and on the right of the QTL peak and their support intervals.

| Trait^a^ | QTL | Linkage  group | QTL  peak  (cM) | Support  interval  (cM) | QTL peak | | | | Support interval | | | |
| --- | --- | --- | --- | --- | --- | --- | --- | --- | --- | --- | --- | --- |
|  |  |  |  |  | Left marker | | Right marker | | Left marker | | Right marker | |
|  |  |  |  |  | solcap_snp | ST4.03 | solcap_snp | ST4.03 | solcap_snp | ST4.03 | solcap_snp | ST4.03 |
| PY06 | 1 | 5 | 32 | 13-42 | c2_50305 | 5,052,466 | c1_14801 | 5,053,838 | c2_11737 | 2,068,427 | c1_5836 | 8,809,052 |
| PY08 | 1 | 5 | 46 | 0-53 | c2_43535 | 9,273,361 | c2_55894 | 9,558,289 | c2_23776 | 65,193 | c1_15638 | 13,087,731 |
| FM07 | 1 | 5 | 26 | 18-29 | c2_11829 | 4,041,510 | c2_22986 | 4,279,075 | c2_11685 | 2,288,291 | c2_23055 | 4,936,332 |
|  | 2 | 7 | 66 | 52-70 | c1_10020 | 45,145,834 | c2_33495 | 45,145,932 | c1_7515 | 41,721,163 | c2_45188 | 46,763,232 |
| FM08 | 1 | 1 | 9 | 0-32 | c2_6713 | 2,068,305 | c2_21097 | 2,589,277 | c2_51460 | 151,047 | c2_27877 | 6,071,374 |
|  | 2 | 5 | 27 | 0-41 | c2_22959 | 4,434,048 | c2_23052 | 4,906,728 | c2_23776 | 65,193 | c1_5836 | 8,809,052 |
|  | 3 | 7 | 65 | 35-71 | c2_44105 | 44,499,261 | c2_44120 | 44,687,759 | c2_52374 | 5,791,166 | c2_45182 | 46,813,301 |
| FM14 | 1 | 5 | 18 | 0-29 | c2_11685 | 2,288,291 | c1_3840 | 3,134,967 | c2_23776 | 65,193 | c2_23055 | 4,936,332 |
|  | 2 | 7 | 66 | 54-70 | c1_10020 | 45,145,834 | c2_33495 | 45,145,932 | c2_23347 | 42,120,645 | c2_45188 | 46,763,232 |
| SG06 | 1 | 2 | 97 | 34-117 | c2_22890 | 42,663,931 | c2_22853 | 42,733,475 | c1_11494 | 26,262,335 | c2_24869 | 46,387,242 |
|  | 2 | 3 | 44 | 31-50 | c2_36469 | 38,175,099 | c1_10879 | 38,177,443 | c1_16267 | 17,239,217 | c1_10514 | 40,768,867 |
| SG07 | 1 | 5 | 19 | 4-32 | c1_3840 | 3,134,967 | c1_3803 | 3,585,641 | c2_23846 | 727,316 | c1_14801 | 5,053,838 |
| SG08 | 1 | 5 | 28 | 15-32 | c2_22959 | 4,434,048 | c2_23052 | 4,906,728 | c2_11712 | 2,149,871 | c1_14801 | 5,053,838 |
| *SG07 | 1 | 2 | 104 | 68-125 | c2_27271 | 44,458,744 | c1_964 | 44,471,630 | c1_9944 | 34,461,501 | c1_11458 | 47,625,735 |
| *SG08 | 1 | 2 | 115 | 60-124 | c1_7872 | 46,195,203 | c1_7871 | 46,195,322 | c2_39178 | 32,148,414 | c1_11458 | 47,625,735 |
| *DM07 | 1 | 2 | 115 | 63-121 | c1_7872 | 46,195,203 | c1_7871 | 46,195,322 | c2_13060 | 32,996,497 | c2_35687 | 47,324,196 |
|  | 2 | 6 | 36 | 31-73 | c2_3962 | 4,889,182 | c2_52241 | 5,790,649 | c2_39216 | 2,792,921 | c2_19556 | 48,069,790 |
| ST08 | 1 | 4 | 2 | 0-4 | c1_7574 | 117,620 | c2_23593 | 635,886 | c1_7574 | 117,620 | c2_23600 | 656,192 |
| *ST08 | 1 | 4 | 2 | 0-4 | c1_7574 | 117,620 | c2_23593 | 635,886 | c1_7574 | 117,620 | c2_23600 | 656,192 |
| NS06 | 1 | 1 | 0 | 0-52 | c2_51460 | 151,047 | c2_51460 | 151,047 | c2_51460 | 151,047 | c2_55008 | 12,994,277 |
| NS08 | 1 | 5 | 26 | 12-32 | c2_11829 | 4,041,510 | c2_22986 | 4,279,075 | c2_52084 | 1,902,218 | c1_14801 | 5,053,838 |
| NI06 | 1 | 1 | 0 | 0-69 | c2_51460 | 151,047 | c2_51460 | 151,047 | c2_51460 | 151,047 | c2_2721 | 58,197,448 |
| NI07 | 1 | 1 | 0 | 0-8 | c2_51460 | 151,047 | c2_51460 | 151,047 | c2_51460 | 151,047 | c2_6906 | 1,606,674 |
|  | 2 | 5 | 25 | 8-34 | c2_11829 | 4,041,510 | c2_22986 | 4,279,075 | c2_33543 | 1,505,540 | c2_52436 | 6,147,926 |
| NI08 | 1 | 5 | 27 | 12-32 | c2_22959 | 4,434,048 | c2_23052 | 4,906,728 | c2_52084 | 1,902,218 | c1_14801 | 5,053,838 |

*Maturity-corrected phenotypes.

^a^Traits: plant yield (PY), foliage maturity (FM), specific gravity (SG), skin texture (ST), and internal heat necrosis severity (NS) and intensity (NI).

**Table S4** Fixed-effect interval mapping (FEIM) for B2721 traits evaluated over four years (2006-8 and 2014).

| Trait^a^ | QTL | Linkage  group | Position  (cM) | Left marker | | Right marker | | LRT^b^ | LOD^c^ | $R_{\mathrm{adj}}^{2}$^d^  (%) |
| --- | --- | --- | --- | --- | --- | --- | --- | --- | --- | --- |
|  |  |  |  | solcap_snp | ST4.03 | solcap_snp | ST4.03 |  |  |  |
| PY08 | 1 | 5 | 45 | c2_43535 | 9,273,361 | c2_55894 | 9,558,289 | 39.05 | 8.48 | 20.5 |
| FM07 | 1 | 5 | 26 | c2_11829 | 4,041,510 | c2_22986 | 4,279,075 | 93.00 | 20.19 | 46.0 |
|  | 2 | 7 | 66 | c1_10020 | 45,145,834 | c2_33495 | 45,145,932 | 35.01 | 7.60 | 18.5 |
| FM08 | 1 | 5 | 28 | c2_22959 | 4,434,048 | c2_23052 | 4,906,728 | 113.12 | 24.56 | 52.7 |
| FM14 | 1 | 5 | 26 | c2_11829 | 4,041,510 | c2_22986 | 4,279,075 | 56.34 | 12.24 | 30.1 |
|  | 2 | 7 | 66 | c1_10020 | 45,145,834 | c2_33495 | 45,145,932 | 29.63 | 6.43 | 15.4 |
| SG06 | 1 | 2 | 42 | c2_41963 | 27,511,649 | c2_41975 | 27,547,993 | 26.67 | 5.79 | 13.4 |
|  | 2 | 8 | 90 | c2_7353 | 48,895,181 | c1_13116 | 49,071,663 | 28.72 | 6.24 | 14.6 |
|  | 3 | 10 | 65 | c2_27827 | 50,697,563 | c2_27829 | 50,782,097 | 29.35 | 6.37 | 15.0 |
| SG07 | 1 | 5 | 21 | c1_3840 | 3,134,967 | c1_3803 | 3,585,641 | 31.48 | 6.83 | 16.4 |
| SG08 | 1 | 5 | 29 | c2_23052 | 4,906,728 | c2_23055 | 4,936,332 | 57.98 | 12.59 | 30.4 |
| *SG07 | 1 | 2 | 105 | c2_27271 | 44,458,744 | c1_964 | 44,471,630 | 27.00 | 5.86 | 13.1 |
| *SG08 | 1 | 2 | 105 | c2_27271 | 44,458,744 | c1_964 | 44,471,630 | 27.60 | 5.99 | 13.4 |
|  | 2 | 5 | 100 | c2_8460 | 50,479,738 | c2_8507 | 50,583,686 | 27.05 | 5.87 | 13.1 |
| *DM07 | 1 | 6 | 36 | c2_24297 | 35,798,871 | c2_24322 | 36,214,491 | 30.44 | 6.61 | 15.0 |
| *DM08 | 1 | 11 | 37 | c2_2896 | 7,479,525 | c2_12368 | 8,096,420 | 27.34 | 5.94 | 13.2 |
| ST08 | 1 | 4 | 2 | c1_7574 | 117,620 | c2_23593 | 635,886 | 56.21 | 12.21 | 29.5 |
|  | 2 | 9 | 92 | c2_14640 | 50,367,678 | c2_14641 | 50,367,972 | 27.13 | 5.89 | 13.6 |
| *ST08 | 1 | 4 | 2 | c1_7574 | 117,620 | c2_23593 | 635,886 | 57.21 | 12.42 | 29.0 |
|  | 2 | 9 | 108 | c2_27054 | 52,799,014 | c2_27003 | 52,996,314 | 28.92 | 6.28 | 14.2 |
| NS06 | 1 | 1 | 2 | c2_36664 | 535,454 | c2_36668 | 559,640 | 26.02 | 5.65 | 13.0 |
| NS08 | 1 | 5 | 27 | c2_22959 | 4,434,048 | c2_23052 | 4,906,728 | 26.45 | 5.74 | 13.2 |
| NI08 | 1 | 5 | 28 | c2_22959 | 4,434,048 | c2_23052 | 4,906,728 | 30.37 | 6.60 | 15.6 |
| NI14 | 1 | 5 | 27 | c2_22959 | 4,434,048 | c2_23052 | 4,906,728 | 26.42 | 5.74 | 13.5 |
| *NI14 | 1 | 4 | 123 | c2_34890 | 67,953,469 | c2_34875 | 67,981,117 | 27.43 | 5.96 | 13.6 |

*Maturity-corrected phenotypes.

^a^Traits: plant yield (PY), foliage maturity (FM), specific gravity (SG), skin texture (ST), and internal heat necrosis severity (NS) and intensity (NI).

^b^Likelihood ratio test (LRT).

^c^Logarithm of the odds (LOD).

^d^Adjusted R-squared ($R_{\mathrm{adj}}^{2}$) in percentage.
